# Estimation of splicing metrics for NMD-sensitive transcripts

**DOI:** 10.64898/2026.06.30.735642

**Authors:** Lev Zavileyskiy, Maria Vlasenok, Antonina Kuznetsova, Dmitry A. Skvortsov, Dmitri D. Pervouchine

## Abstract

Alternative splicing is commonly quantified using the Percent-Spliced-In (PSI) metric, which measures the relative abundances of alternatively spliced isoforms. However, some transcript isoforms are targeted by the nonsense-mediated mRNA decay (NMD) pathway, introducing a strong bias that leads to underestimation of their true splicing rates. To correct for this bias, we developed an analytical framework and a set of statistical models employing a linear fractional transformation that depends on a single parameter to capture the degradation rate of transcripts annotated as NMD targets relative to normal mRNA decay. Using Gaussian mixture models, we demonstrated a clear separation of splicing events into two classes, NMD-sensitive and NMD-evading, with the former exhibiting strong upregulation upon NMD inhibition and the latter showing little or no response. Moreover, NMD-evading events displayed higher coding potential and stronger translation signals both upstream and downstream of the stop codon, which are characteristic of NMD escape through translational readthrough. We further showed that incorporation of event-specific relative decay rates improves the interpretation of differential splicing patterns for transcripts annotated as NMD targets. In sum, our results provide a solid framework for corrected estimation of splicing metrics in NMD-targeted transcripts from short-read RNA-seq data.

## Introduction

Splicing is a key post-transcriptional pre-mRNA processing step in which introns are removed and exons are joined together to produce mature transcripts [1]. In higher eukaryotes, more than 90% of multi-exon genes undergo alternative splicing (AS) generating multiple mRNA isoforms from a single gene, which enables tissue-specific regulation and environmental adaptation [2, 3]. Translation of alternatively spliced isoforms substantially expands proteome diversity, although the extent and importance of this contribution remain a subject of ongoing debate [4, 5, 6].

Quantification of AS can be performed using various approaches, including methods based on short read RNA sequencing (RNA-seq) [7, 8, 9, 10]. The most widely used metric, the Percent Spliced-In (PSI or Ψ) ratio, estimates the abundance of transcript isoforms containing an alternatively spliced exon relative to the total abundance of inclusion and skipping isoforms [11]. Originally formulated for cassette exons, this definition also applies to more complex types of AS events [12]. In RNA-seq data, Ψ is typically estimated as the fraction of reads supporting exon inclusion among all reads supporting either inclusion or skipping. Although several refinements to this definition have been proposed to account for peculiarities of the RNA-seq protocol, such as PCR duplicates and non-unique read mappings [13, 9], quantification of AS by the Ψ metric data remains a golden standard in most current transcriptomic studies.

Apart from technical biases inherent to short-read RNA-seq protocols, a source of systematic error in estimating Ψ arises from the fact that a fraction of transcript isoforms generated by AS are degraded by the nonsense-mediated mRNA decay (NMD) pathway, which selectively eliminates transcripts harboring premature termination codons (PTCs) [14, 15]. Since most RNA-seq protocols employ poly(A) selection to increase productivity, the majority of short reads used to compute the observable Ψ metric originate from the cytosolic RNA fraction and reflect the equilibrium concentrations of transcripts after a fraction of them has been degraded by NMD. In contrast, the actual splicing rate *r*, reflecting transcript abundances in the nucleus where AS occurs prior to NMD-mediated mRNA degradation, cannot be measured directly and may differ from the observed Ψ metric. This discrepancy is particularly relevant for transcript isoforms predicted or annotated as NMD targets, whose observed cytoplasmic abundances may substantially underestimate the actual splicing rates.

The classic NMD model posits that the premature status of a stop codon is recognized through a translation-dependent mechanism in which an early terminating ribosome fails to displace downstream exon-junction complexes (EJCs) [16]. This configuration promotes activation of the ATP-dependent RNA helicase UPF1, which interacts with UPF2, UPF3, and the SMG1 kinase, leading to UPF1 phosphorylation and recruitment of downstream decay factors [17]. These include SMG6, an endonuclease that promotes direct cleavage of NMD-targeted transcripts, and SMG7, an adaptor protein that recruits factors involved in deadenylation, decapping, and exonucleolytic degradation [18]. The corresponding annotation rule states that a PTC is more likely to trigger NMD when it is located approximately 50–55 nucleotides upstream of an EJC (the 50-nt rule), although NMD efficiency is also influenced by the position of the PTC relative to the start codon, exon lengths, and proximity to the 3’-untranslated region (3’-UTR) [19].

The coupling between AS and NMD, termed AS-NMD or unproductive splicing, represents a widespread mechanism of post-transcriptional gene regulation [20, 21, 22]. Through regulated splicing events, this mechanism generates transcripts containing PTCs, which are subsequently targeted for NMD-mediated degradation. AS-NMD forms complex autoregulatory and cross-regulatory feedback loops in which splicing factors regulate their own expression and that of other RNA-binding proteins, thereby shaping gene expression across regulatory networks [23, 22, 24]. Estimating the splicing rate *r* upstream of NMD-mediated degradation is therefore essential for accurately characterizing such networks, as rapid degradation of NMD-sensitive isoforms can mask underlying regulatory interactions when NMD is intact.

In this work, we developed a statistical model based on a simple kinetic framework to reconstruct true exon inclusion rates *r* from observed Ψ values. The model describes the relationship between Ψ and *r* using a family of linear fractional transformations that depend on a single parameter *γ*, which captures the relative degradation rate of NMD-targeted transcripts relative to the natural decay rates of protein-coding mRNAs. Parameters of these models can be inferred from RNA-seq experiments that enable accurate measurement of *r* and Ψ distinguishing nuclear splicing outcomes from the effects of cytoplasmic RNA surveillance. Among them, experiments performed with and without NMD inactivation provide a particularly convenient method, yielding paired measurements of Ψ and *r* under untreated and NMD-inhibited conditions, respectively.

We first estimate *γ* directly from NMD inhibition experiments, focusing primarily on the co-depletion of SMG6 and SMG7, which strongly suppresses NMD by simultaneously inhibiting its two major degradation branches [25]. We then extend the approach by applying a two-component Gaussian mixture model, revealing that transcripts annotated as NMD targets are not uniformly affected by NMD. Instead, they segregate into two groups, NMD-sensitive and NMD-evading, with two distinct average *γ* values. Applying the same model to other relevant datasets, including nuclear–cytosolic RNA fractionation and cycloheximide-mediated NMD inhibition experiments, we find that this classification is largely consistent across different conditions. We further show that NMD-evading events tend to have higher coding potential and stronger Ribo-seq signatures, suggesting NMD escape via translational readthrough in a subset of events. Next, we demonstrate that event-specific relative decay rates improve the interpretation of the relationship between differential splicing and gene expression. Finally, we validate six predicted NMD-sensitive events in human PC3 prostate adenocarcinoma cells and show that one of them (*NFAT5*) behaves as NMD-evading, possibly via translational readthrough.

## Methods

### Cell cultures

Human PC3 prostate adenocarcinoma cells were maintained in Dulbecco’s Modified Eagle medium / Nutrient Mixture F-12 with 10% v/v fetal bovine serum (FBS), 1% GlutaMAX, 0.05 mg/ml streptomycin, and 50 units/ml penicillin (all products from Thermo Fisher Scientific). Cells were maintained at 37°C in 5% CO_2_ and plated at a density of 120,000 cells per well in a 12-well plate. To inhibit NMD, cycloheximide (CHX) was added to the cells 3 hours before harvest giving a final concentration of 300 mg/ml. All experiments were performed with at least three independent biological replicates.

### RNA purification and cDNA synthesis

Total RNA was isolated using the guanidinium thiocyanate–phenol–chloroform method with ExtractRNA Reagent (Evrogen), following the manufacturer’s protocol. To eliminate residual genomic DNA, 1 microgram of total RNA was treated with RNase-free DNase I (Thermo Fisher Scientific) at 37^◦^C for 30 min. Subsequently, 500 ng of DNase-treated RNA was used for complementary DNA (cDNA) synthesis using Magnus First Strand cDNA Synthesis Kit for RT-qPCR (Evrogene) to a final volume of 20 µl. cDNA was diluted 1:5 with nuclease-free water for quantitative PCR (qPCR).

### qPCR

qPCR reactions were performed in triplicate in a final volume of 12 µl using 96-well plates. Each reaction contained 420 nM gene-specific primers, 2 µl of cDNA, and 5XqPCRmix-HS SYBR reaction mix (Evrogen). Primer sequences used for qPCR are listed in Table S1. A sample without reverse transcriptase enzyme was included as a control to verify the absence of genomic DNA contamination. Amplification was carried out on CFX96 Real-Time System (Bio-Rad), with the following parameters: 95°C for 5 min, followed by 39 cycles at 95°C for 20 s, 58°C for 20 s and 72°C for 20 s, ending at 72°C for 5 min. Primer amplification efficiencies were determined for each primer pair using standard calibration curves. Relative isoform expression levels were calculated using the efficiency-corrected method, with all primer pairs exhibiting amplification efficiencies greater than 90%.

### RNA sequencing data

The poly(A)^+^ RNA-seq datasets from HeLa cells subjected to NMD factor knockdowns (KD) or cycloheximide treatment [25, 26], as well as nuclear and cytosolic RNA fractionation experiments performed in K562 and HepG2 cells [27, 28], were obtained from the SRA repository in FASTQ format. The UPF1 knockdown RNA-seq data from HCT116 cells were downloaded from the E-MTAB-13788 dataset in BAM format [29]. Sample IDs for these RNA-seq datasets are listed in Table S2. The poly(A)^+^ RNA-seq data of human tissues (Table S3) from the GTEx project V7 were downloaded from dbGaP in FASTQ format, retaining only samples with read length of 75 nt and at least 20 million short reads. The poly(A)^+^ RNA-seq datasets of K562 and HepG2 cells subjected to RBP knockdowns (469 experiments, two bioreplicates each) were obtained from the ENCODE consortium [30] in BAM format (alignments to GRCh38). Perturbations with less than twofold inactivation of the target RBP or datasets lacking separation of control and perturbed samples by hierarchical clustering of DESeq2-normalized gene counts were excluded. FASTQ reads were aligned to the GRCh38 human genome using STAR v2.7.8a [31] with GENCODE v43 annotation as a reference [32]. Gene counts were calculated using FeatureCounts utility [33]. Differential expression analysis was performed using the DESeq2 package [34].

### AS event catalog and quantification

A catalog of AS events producing NMD isoforms was constructed from Ensembl v108 [35] and CHESS 3 transcriptome annotations [36] using the NMDj pipeline [37]. This pipeline identifies characteristic introns that distinguish protein-coding from NMD isoforms and assigns each intron a weight based on the number of isoforms in which it occurs. Before running NMDj, CHESS gene identifiers were matched to Ensembl gene identifiers to remove duplicate entries. NMD transcripts from Ensembl were selected on the basis of transcript biotype attribute excluding transcripts lacking annotated start or stop codons. NMD transcripts from CHESS were predicted by formal translation using the 50-nt rule. The resulting lists were merged, excluding CHESS events that shared characteristic introns with any Ensembl event from the same gene.

This procedure yielded a catalog of 14,558 AS events predicted to generate NMD isoforms, of which 3,923 met the gene expression and read-coverage criteria in the cell lines analyzed in this study (see below). From this catalog, we selected 1,163 poison exon (PE) and essential exon (EE) events, which generate NMD isoforms upon exon inclusion and skipping, respectively, and a manually curated subset of 32 AS events producing experimentally validated NMD isoforms based on a literature search (Table S4). A control set of 3,756 AS events not producing NMD isoforms was extracted from Ensembl v108 and CHESS annotations by selecting cassette (skipped) exons, as described previously [38].

Split-read counts supporting splice junctions were extracted from RNA-seq alignments using the IPSA pipeline with the default settings [12]. For AS events predicted to generate NMD isoforms, the splicing ratio was calculated as Ψ = *a*/(*a* + *b*), where *a* and *b* denote the weighted numbers of reads supporting the NMD and protein-coding isoforms, respectively. For AS events producing only protein-coding isoforms, Ψ was calculated with respect to the inclusion isoform. AS events with ΔΨ = 0 or the average denominator of the Ψ ratio below 15 reads were excluded, following previously described filtering criteria [39].

A schematic overview of the analysis workflow, including the catalog construction, extraction of split-read counts, estimation of event-specific transcript decay rates, and Gaussian mixture models fitting (see Results section), is provided in Figure S1. It is implemented as a Python script available at the Github repository [40] to enable reproducible analysis.

### Stop codon readthrough and coding potential

Ribosome profiling data from human HCT116 cells subjected to NMD inhibition were obtained from BioStudies (E-MTAB-13837) in BAM format following initial read processing and quality control [29]. In brief, short reads were demultiplexed and adapter-trimmed, retaining only reads 16–40 nt in length. The remaining reads were aligned to the GRCh38 reference genome with STAR v2.7.10b using GENCODE v42 supplemented with SIRVome/ERCC annotations excluding mappings to non-coding RNAs [29]. BAM alignments were converted to BigWig format using the bamCoverage utility with 10-nt bins and BPM (bins per million mapped reads) normalization [41]. The sample UPF1 12h 3 was excluded due to a divergent read coverage profile. Only uniquely mapped reads (MAPQ 255) were retained for subsequent analysis.

Ribosome footprint coverage was quantified in regions upstream and downstream of the stop codon in PTC-containing poison exons (*n* = 539), excluding 25-nt window centered on the PTC to avoid the termination-associated ribosome peak. NMD-sensitive and NMD-evading exons were matched by their mean ribosome footprint coverage (*n_S_* = 133 and *n_E_* = 151, respectively). For each exon, the readthrough fraction was calculated as the mean downstream ribosome footprint coverage divided by the mean upstream coverage, considering only regions longer than 5 nt. To exclude the possibility that reads originating from overlapping protein-coding transcripts were misassigned to NMD isoforms, we intersected poison exons with annotated Ensembl CDSs and excluded events with such overlaps.

To assess the coding potential, basewise PhyloP conservation scores across 30 species (27 primates and three non-primate mammals) for the human GRCh38 genome assembly were downloaded from the UCSC Genome Browser [42]. For each PTC-containing poison exon (*n* = 539), the coding potential (ΔPhyloP) was calculated as the difference between the mean PhyloP score at the first and second codon positions and the PhyloP score at the third position, averaged across codons. As described previously [38], protein-coding regions exhibit higher average ΔPhyloP values than non-coding regions due to the degeneracy of the genetic code.

### Statistical analysis

Statistical analyses were performed in Python 3.10. Linear models used to estimate *γ* were fitted by orthogonal distance regression (total least squares) implemented in the scipy.odr library. In all cases, the intercepts were fixed according to the corresponding theoretical model. The stability of *γ* estimates was assessed by bootstrapping using 50 bootstrap samples with replacement, each containing 50% of the events in the corresponding dataset. In Gaussian mixture modelling, model parameters were estimated by maximum likelihood using the optimization routines implemented in the scipy.optimize package. One-sided Mann–Whitney test with continuity correction was used for pairwise comparisons, except for the analyses shown in Figure 5, where a paired *t*-test was used since the data did not depart significantly from normality. Group comparisons in Figures 4 and 5 were corrected for multiple testing using the Bonferroni method. Boxplot whiskers correspond to observations within 1.5 interquartile ranges from the median; outliers were omitted from graphical displays. Statistical tests and significance annotations were performed using the statannotations package. Significance levels of 5%, 1%, 0.1%, and 0.01% are denoted by *, **, ***, and ****, respectively.

## Results

### The kinetic model

We propose a minimal kinetic model (Figure 1A) that links the actual nuclear splicing rate *r* prior to NMD-mediated mRNA degradation to the observed equilibrium Ψ value after partial transcript decay. The model assumes that transcription produces pre-mRNAs from a given gene at a constant rate *v*. In turn, pre-mRNAs are assumed to be spliced co-transcriptionally into either a protein-coding isoform (PC) or a NMD-target isoform (NMD) with probabilities 1 − *r* and *r* (0 ⩽ *r* ⩽ 1) resulting in their effective production rates of (1 − *r*)*v* and *rv*, respectively. As in previous works [39], Ψ and *r* are always defined relative to the NMD isoform, such that higher values indicate a greater abundance of NMD targets.

**Figure 1:**
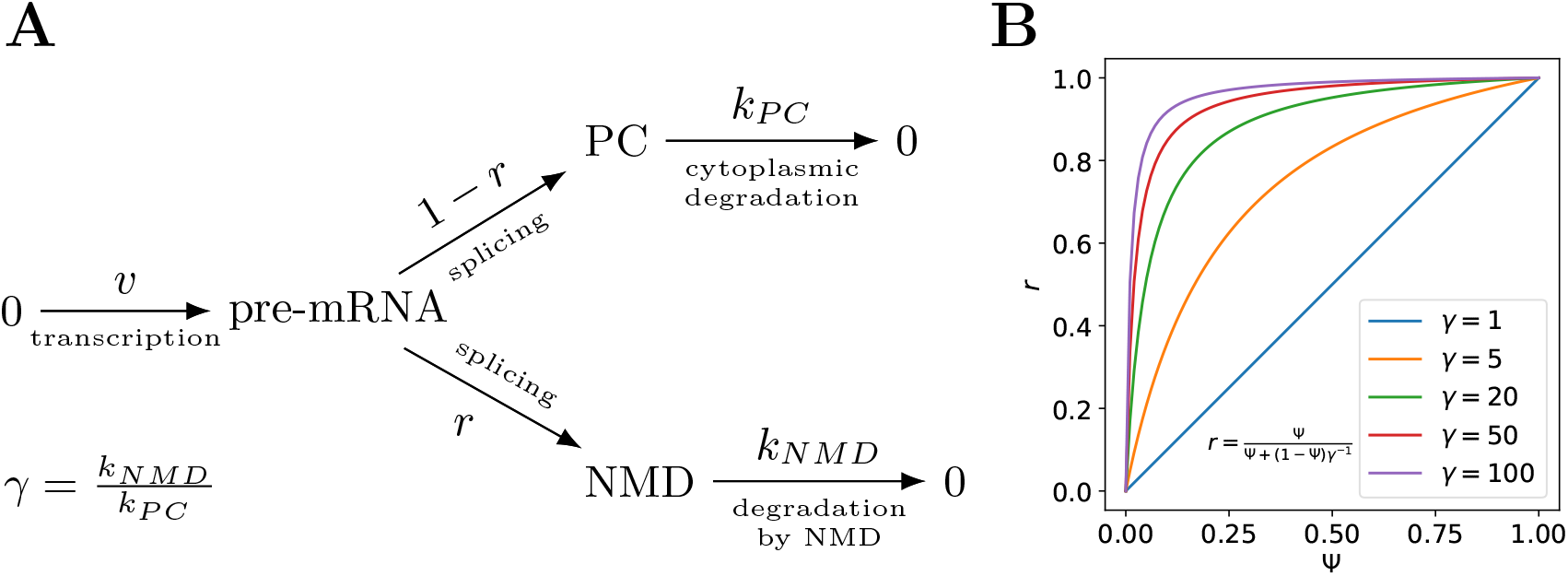
A kinetic model of splicing coupled with NMD. **(A)** Pre-mRNA is transcribed at rate *v* and spliced co-transcriptionally (instantly) into either the protein-coding (PC) isoform, with probability 1 − *r*, or the NMD-targeted (NMD) isoform, with probability *r*. The two isoforms decay at rates *k_PC_* and *k_NMD_*, respectively. **(B)** The relationship between the nuclear splicing rate *r* and the observed Percent Spliced-In value Ψ.

The PC and NMD isoforms are degraded with kinetic rate constants *k_PC_* and *k_NMD_*, respectively, where *k_NMD_* > *k_PC_*. Their ratio, 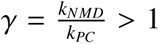, reflects the activity of the NMD pathway toward NMD isoforms, where larger *γ* values correspond to their faster degradation relative to protein-coding isoforms.

The dynamics of PC and NMD isoform concentrations can be described by the following system of ordinary differential equations:

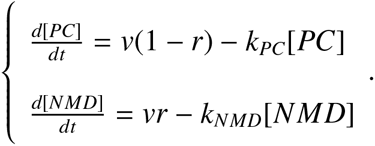

At equilibrium, the concentrations of the PC and NMD isoforms are given by the steady-state conditions:

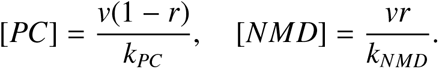

Then, the observed Ψ value can be computed as the ratio of the concentration of the NMD isoform to the combined concentration of NMD and PC isoforms:

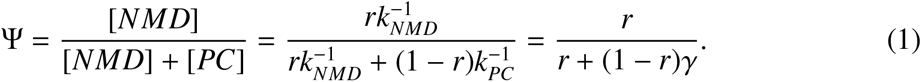

That is, Ψ and *r* are related by a linear fractional transformation Ψ = Ψ(*r*), which satisfies Ψ(0) = 0 and Ψ(1) = 1. The inverse function *r* = *r*(Ψ) is given by a similar expression:

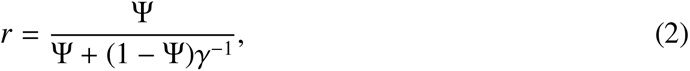

in which the roles of the NMD and PC isoforms are interchanged, and *γ* is replaced by *γ*^−1^.

The family of functions defined by Eq. 2 also satisfies *r*(0) = 0 and *r*(1) = 1 and deviates from the identity function at *γ* = 1, becoming increasingly nonlinear as *γ* increases (Figure 1B). This indicates that the true splicing rate *r* can be substantially larger than the observed Ψ value, especially when Ψ is small. For comparison, if the NMD isoform degrades four times faster than the PC isoform (*γ* = 4), the observed value of Ψ = 0.2 corresponds to *r* = 0.5.

### Direct estimation of relative transcript decay rates

NMD inactivation experiments provide a powerful method for obtaining paired measurements of Ψ and *r* because the value Ψ*_KD_*, that is, Ψ measured under conditions in which the NMD system is inactive and no differential transcript degradation occurs (*γ* = 1), must be equal to the true splicing rate, i.e., Ψ*_KD_*≃*r*.

First, we analyzed RNA-seq datasets obtained upon experimental inactivation of the NMD pathway in HeLa cells, in which a double knockdown of the essential NMD factors SMG6 and SMG7 was performed, resulting in a strong suppression of the NMD pathway [25]. We compared Ψ under control conditions with Ψ*_KD_*≃*r* measured after SMG6/SMG7 double knockdown across three sets of AS events that generate NMD isoforms, among those detectable in the cell lines analyzed: 32 experimentally validated events (VAL, see Table S4), 1,163 annotated cassette exons classified as poison or essential exons (PE+EE), and a broader set of 3,923 complex AS events (ALL) predicted to generate NMD isoforms (see Methods).

To estimate *γ*, we employed three approaches. First, we rewrote Eq. 1 as(Ψ^−1^ – 1) = *γ* (*r*^−1^ – 1), which allows *γ* to be estimated as the slope of a linear regression of Ψ^−1^ − 1 on *r*^−1^ − 1 with fixed intercept. Because this approach becomes unstable when Ψ and *r* are close to zero, it was applied only to observations with Ψ ⩾ 0.01 and *r* ⩾ 0.01. Second, we directly fitted the linear fractional function given by Eq. 2 using the nonlinear least squares method. Finally, we estimated *γ* from the derivative of the function *r* = *r*(Ψ), exploiting the fact that *r*^′^(0) = *γ*. In contrast to the first approach, this method is applicable only to observations near the origin in the (Ψ, *r*) plane. Accordingly, it was estimated by fitting a linear regression with zero-fixed intercept to observations with Ψ < 0.05.

Although the three approaches consistently confirmed selective degradation of NMD isoforms, the estimated values of *γ* varied substantially across methods and event sets. While on the set of validated events all three methods converged to *γ* ≈ 18 (Figure 2A), a significant discrepancy between the regression and derivative-at-zero models was observed for both annotated poison and essential exons (Figure 2B) as well as for the full set of AS events producing NMD isoforms (Figure 2C). Bootstrapping showed that estimates obtained by linearization and nonlinear curve fitting ranged from approximately 2 to 20, depending on the subset of events analyzed, whereas the derivative-at-zero approach yielded consistent estimates *γ* ≈ 18 across all datasets (Figure 2D).

**Figure 2:**
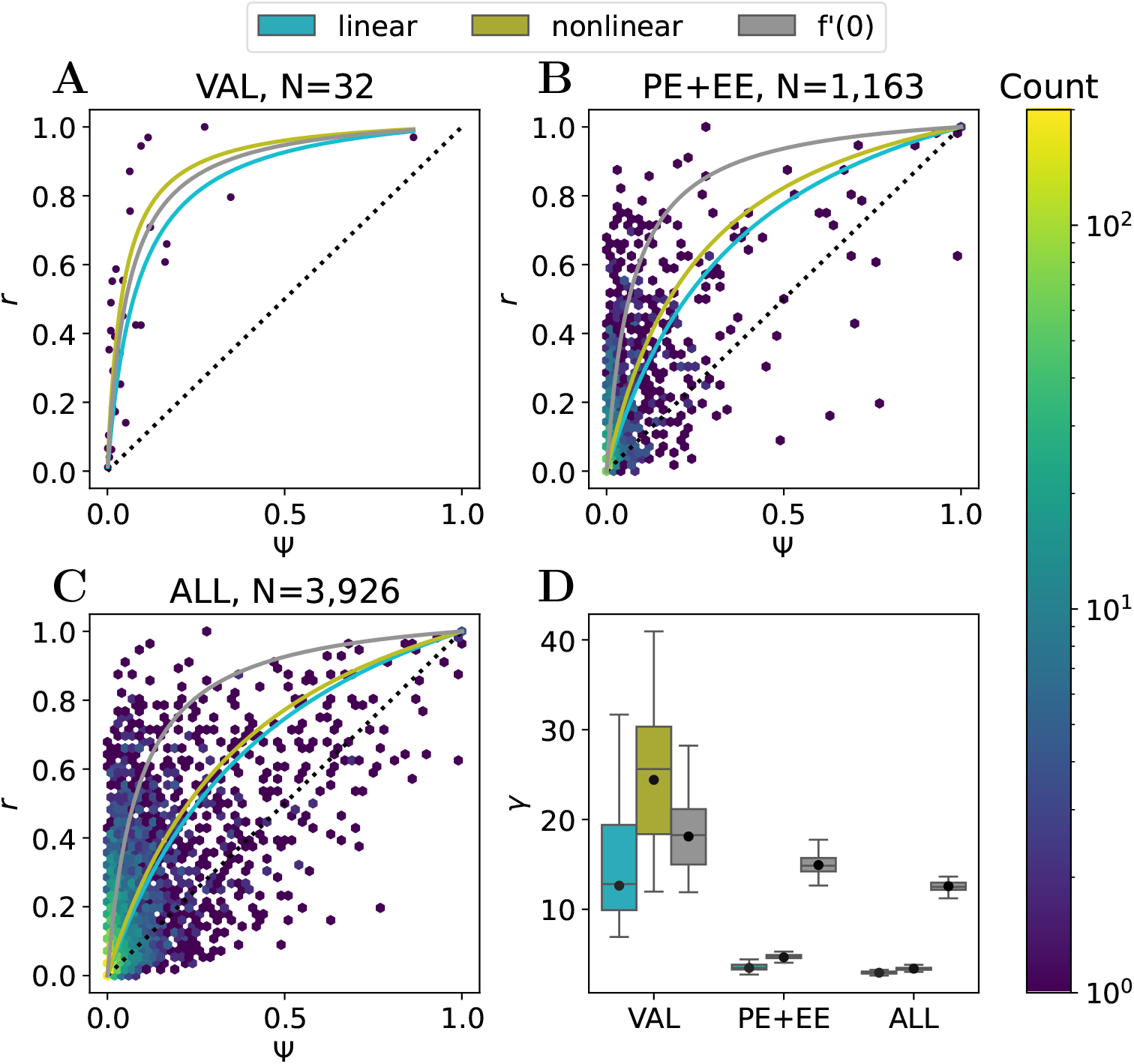
Estimation of the average relative decay rate *γ* from NMD inactivation experiments. **(A–C)** Scatter plots relating splicing metric values measured under unperturbed (Ψ) and NMD-inactivated (*r*) conditions (SMG6/SMG7 codepletion) for different subsets of splicing events: experimentally validated events (VAL), annotated poison and essential exons (PE+EE), and all NMD-generating AS events (ALL). *Linear* denotes linear regression in the transformed coordinates Ψ^−1^ − 1 = *γ*(*r*^−1^ − 1) for Ψ ⩾ 0.05 using events with *r* ⩾ 0.05. *Nonlinear* denotes direct nonlinear least-squares fitting. *f* ^′^(0) denotes estimation from the slope at the origin for events with Ψ_ctrl_ < 0.05. **(D)** Estimates of *γ* obtained from bootstrap samples (*N* = 50), each containing a random half of the events (see Methods).

To a large extent, the observed discrepancies reflect the fact that different models estimate *γ* over different ranges of Ψ and *r*: the linearized model primarily incorporates points with larger values, whereas the derivative-at-zero model instead focuses on small values. Notwithstanding, in all cases a large fraction of events deviated substantially from the theoretical curve expected under a single global *γ* value, with some exhibiting a strong shift toward NMD-mediated mRNA degradation, while others remaining close to the diagonal and showing weak or absent response to NMD inhibition. Moreover, even among apparently responsive events, the spread around the fitted curves was considerably large. All these observations suggest that NMD-generating AS events cannot be adequately described by a model with a single universal degradation parameter.

### Mixture model identifies NMD-responsive splicing events

We next employed a statistical model explicitly accounting for heterogeneity in NMD sensitivities across AS events that does not require a fixed value of *γ*. The fact that a fraction of AS events annotated as producing NMD isoforms clustered around the diagonal (Figure 2C) shows that not all transcripts annotated as NMD targets actually respond to NMD inhibition [43, 44, 45].

Accordingly, we hypothesized that the collection of splicing events comprises a mixture of two components: NMD-evading (E), whose NMD transcripts decay at the baseline rate *γ_E_* ≈ 1, and NMD-sensitive (S), whose NMD transcripts decay at a higher relative rate with *γ_S_* > *γ_E_*. Assuming that π and 1 − *π* are the proportions of NMD-sensitive and NMD-evading events, respectively, we modeled the conditional probability density of Ψ given *r* as a mixture of Gaussian distributions:

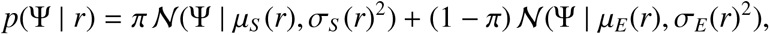

where *N*(Ψ | *µ*, *σ*^2^) denotes normal probability density with the mean *µ* and variance *σ*^2^. The means of these distributions vary as a function of *r* according to Eq. 1 with relative decay rate parameters *γ_S_* and *γ_E_*, respectively:

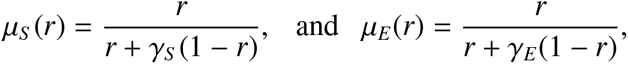

whereas the standard deviations *σ_S_* (*r*) and *σ_E_*(*r*) depend quadratically on *r* in order to account for reduced variation near extreme inclusion levels of 0 and 1. Then the posterior probability of a splicing event being NMD-sensitive is given by

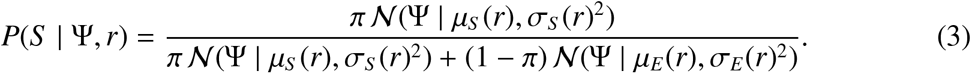

In application to the bivariate data from SMG6/SMG7 double knockdown experiments, we first estimated the baseline parameter *γ_E_* using a set of AS events that do not generate NMD isoforms, and confirmed that the model yields *γ_E_*estimate close to 1 (Figure S2). We then estimated the remaining parameters (*γ_S_*, π, and coefficients of σ*_S_* and σ*_E_*) by maximizing the likelihood of the mixture model on validated, annotated poison and essential exons, and the full set of NMD-generating AS events.

The first model (VAL) yielded *γ_S_* ≈ 16, while the other two models (PE+EE and ALL) converged to a remarkably similar estimate of *γ_S_* ≈ 8, differing only in π, the inferred proportion of NMD-sensitive events, which expectedly decreased from 95% for validated events to 66% for the full set of predicted events (Figure 3A–C). In all cases, AS events assigned higher posterior probabilities of being NMD-sensitive were shifted above the diagonal, while NMD-evading events largely followed the baseline relationship corresponding to weak or absent differential degradation. The proportion of NMD-evading events was the largest for the full set of events, where NMD targets were inferred from transcript annotations based on the 50-nt rule, confirming that it is an imperfect predictor of NMD sensitivity [46].

**Figure 3:**
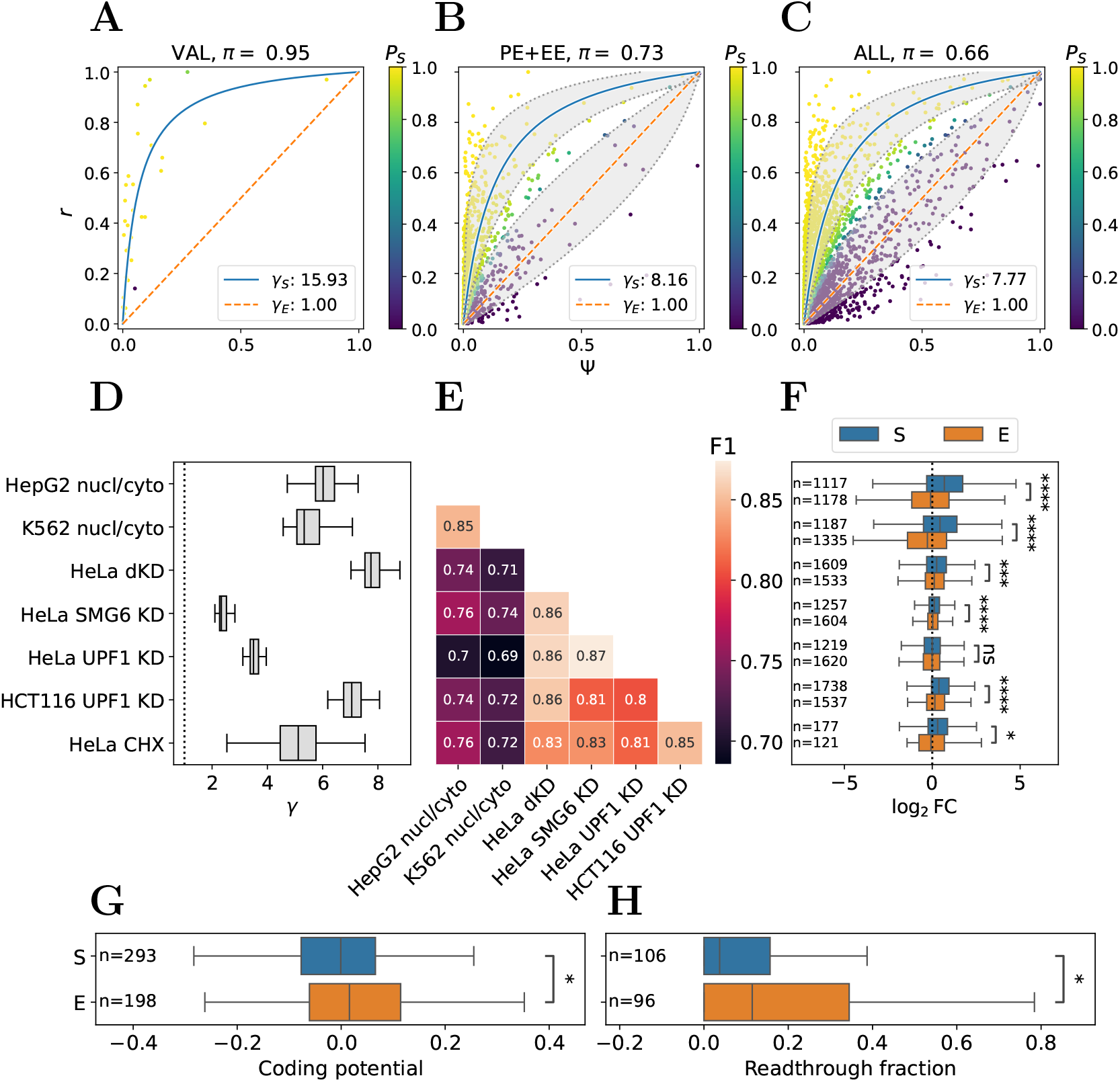
Gaussian mixture model with two components. **(A–C)** Two-component Gaussian mixture model fitted to SMG6/SMG7 codepletion RNA-seq data. Each point represents an AS event and is colored according to the posterior probability of belonging to the NMD-sensitive (S) component. The estimated fraction of NMD-sensitive events, π, is indicated above each plot. Solid and dotted lines correspond to the expected relationships *r* = *r*(Ψ) for *γ_S_* and *γ_E_*, respectively. Shaded areas denote ±1 standard deviation around the mean. The remaining elements of the plots are as in Figure 2A–C. **(D)** Bootstrap estimates (*N* = 50) of *γ_S_*. **(E)** Similarity between NMD-sensitive/NMD-evading classification, measured by pairwise *F*_1_-scores. **(F)** Relative host gene expression changes (log_2_ *FC*) upon NMD inactivation. **(G)** Coding potential upstream of the PTC, defined as the mean PhyloP score in the first two codon positions minus the mean PhyloP score in the third position. **(H)** Readthrough fraction, defined as the ratio of average Ribo-seq coverage downstream versus upstream of the PTC. Asterisks denote significance levels based on Mann–Whitney tests (****, *p* < 0.0001; ***, *p* < 0.001; **, *p* < 0.01; *, *p* < 0.05).

To assess the robustness of the inferred *γ_S_* values across experiments, we applied the mixture model in a bootstrap mode to multiple datasets employing different strategies for paired measurements of Ψ and *r*, including knockdowns of individual NMD factors (UPF1, SMG6), SMG6/SMG7 double knockdown, cycloheximide (CHX) treatment, and nuclear-cytosolic RNA fractionation experiments, with respect to the full set of NMD-generating AS events.

All experimental protocols consistently supported *γ_S_* being greater than one, but its estimated magnitude varied substantially (Figures 3D and S3). The highest values were observed for SMG6/SMG7 double knockdown in HeLa cells, consistent with simultaneous suppression of the two major NMD degradation branches [47]. Notably, UPF1 knockdown in HCT116 cells yielded the second highest estimate, whereas UPF1 knockdown in HeLa cells resulted in a substantially lower *γ_S_*. Nuclear-cytosolic fractionation and CHX-mediated NMD inhibition also yielded relatively high estimates, while individual SMG6 and UPF1 knockdowns in HeLa cells produced lower values than the SMG6/SMG7 double knockdown. Collectively, these results highlight the strong dependence of estimated NMD efficiency on cellular context and experimental conditions, even when targeting the same core NMD factor.

Next, we evaluated the concordance between NMD-sensitive and NMD-evading classifications obtained from different datasets, where a NMD-sensitive event was defined as having *r* > 0.05 and posterior probability *P*(*S* | Ψ, *r*) > 0.5, using pairwise *F*_1_-scores (Figure 3E). All pairwise *F*_1_-scores exceeded 68%, indicating generally consistent classification. To assess the robustness of this result to the choice of the threshold, we considered a range of cutoffs, 0.5 < *t* < 1, where events were classified as NMD-sensitive if *P*(*S* | Ψ, *r*) > *t* and as NMD- evading if *P*(*S* | Ψ, *r*) ⩽ 1 − *t*. Events with intermediate probabilities, 1 − *t* < *P*(*S* | Ψ, *r*) ⩽ *t*, were left unclassified. Increasing classification stringency raised the mean *F*_1_-scores above 90% for comparisons within each experimental group (NMD inhibition or nuclear–cytosolic fractionation), but not for comparisons between the two groups (Figure S4).

Clustering of the similarity matrix revealed that most classification discrepancies arose from differences between NMD inhibition and RNA fractionation experiments (Figures 3E and S5A). In particular, AS events classified as NMD-sensitive in the latter but not in other experiments, were strongly enriched with intron retentions, suggesting that their attribution to NMD-sensitive class may be driven by nuclear enrichment of unspliced transcripts (Figure S5B). Thus, RNA fractionation experiments may introduce a distinct source of variation that is related to incomplete or ongoing post-transcriptional splicing and not to NMD sensitivity.

Consistent with the enhanced degradation of NMD isoforms generated by NMD-sensitive AS events, their host genes showed significantly greater positive expression changes (log *FC*) both in the cognate SMG6/SMG7 co-depletion data and in other experiments (Figure 3F). Although nuclear retention of incompletely spliced transcripts may contribute to their relative nuclear enrichment, it is unlikely to drive the positive shift of log *FC* because the majority of AS events were consistent between RNA fractionation and NMD inactivation experiments (Figure 3E).

One might expect NMD-evading poison exons to have a greater coding potential [48]. To test this hypothesis, we compared the coding potentials of NMD-sensitive and NMD-evading poison exons using ΔPhyloP, defined as the difference between the mean PhyloP scores at the first two codon positions and the PhyloP score at the third position (see Methods). This metric exploits the three-nucleotide periodicity of protein-coding sequences, reflecting the weaker selective constraint generally observed at the third codon position [49, 50, 38]. NMD-evading poison exons exhibited higher coding potential than NMD-sensitive poison exons both upstream (Figure 3G, *P* = 0.019) and downstream of the PTC (*P* = 0.095). These patterns may reflect stronger purifying selection on sequences upstream of the PTC in NMD-evading isoforms and potentially on downstream sequences in a subset of isoforms undergoing translational readthrough.

We next investigated whether NMD-evading events exhibit known features associated with NMD escape. These features include the position of the PTC relative to the translation start site, the location of the PTC within the exon, and the potential for translation reinitiation at a downstream AUG codon [51]. A random forest classifier based on these features distinguished NMD-sensitive from NMD-evading events better than chance alone, but its predictive performance was modest (ROC-AUC = 0.65). Distance between the translation start site and the PTC was the most informative feature, while other features also had positive contributions (Figure S6). These results are consistent with the notion that NMD evasion is influenced by multiple factors such as less efficient NMD at start-proximal PTCs, modulation of the relationship between a PTC and downstream EJCs by exon architecture, and translation reinitiation at downstream AUG codons [52, 19, 53]. Our analysis only identified associations with NMD sensitivity rather than estimating specific contributions of any of these mechanisms.

Finally, we examined whether NMD-evading events were more frequently associated with translational readthrough at the PTC in poison exons. To conduct this analysis, we focused on Ribo-seq data collected under NMD inhibition because NMD-mediated degradation reduces the number of informative ribosome footprints. The readthrough fraction (see Methods) showed that ribosome footprints extended beyond the PTC more frequently in NMD-evading than in NMD-sensitive events (Figure 3H), supporting readthrough as a potential route of NMD escape in a subset of events. To rule out a possibility that this effect was caused by low read coverage, we repeated the analysis requiring more than 5 or more than 10 reads in the region upstream of the PTC and consistently observed elevation of the readthrough fraction in NMD-evading poison exons (Figure S7). Read coverage in samples with intact NMD was insufficient to estimate the readthrough frequency reliably. In sum, the observed patterns support an association between readthrough and NMD evasion, although they do not establish readthrough as the cause of NMD-escape for individual transcripts.

### Event-specific relative decay rates correlate with transcriptional response

The mixture model presented in the previous section showed that global estimates of NMD efficiency can capture the behavior of AS events with two distinct average *γ* values. The biological reality, however, is likely more complex and may involve a continuum of degradation rates rather than two discrete classes. Considering this, we explored the behavior of each AS event *i* characterized by its individual *γ_i_*value, which can be expressed through Ψ*_i_* and *r_i_* directly from Eq. 1 as

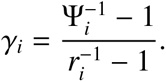

To determine whether event-specific *γ_i_* values (hereafter, *γ* without index *i*) are consistent with the existing measure of NMD efficiency, we compared them with transcript-level NMD response scores reported by Colombo et al. [25]. In that study, NMD efficiency was quantified using a log_2_ fold-change score that integrates expression differences between NMD-factor knockdown and control conditions with those between NMD-factor rescue and knockdown conditions, so that higher positive scores indicate stronger NMD-dependent transcript accumulation. Log-transformed event-specific *γ* values showed significant positive Spearman correlations with the NMD response log_2_ fold-change scores all datasets, with the strongest association observed for the SMG6/SMG7 double knockdown (ρ = 0.44; Figure S8). These results provide independent confirmation that event-specific *γ* values capture biologically meaningful variation in NMD efficiency, despite being inferred from splicing changes rather than directly from transcript abundance.

We next asked whether *γ* values also reflect the strength of coupling between AS and gene expression by analyzing a panel of human tissue transcriptomes from the GTEx project. We expected that events with higher *γ* values would exhibit stronger negative associations between Ψ and host gene expression level (*expr*). Consistent with this expectation, Spearman correlation coefficients of Ψ and *expr* were lower for NMD-sensitive than for NMD-evading events and decreased progressively with increasing *γ* (Figure 4A), indicating that it reflects the efficiency of AS–NMD coupling.

**Figure 4:**
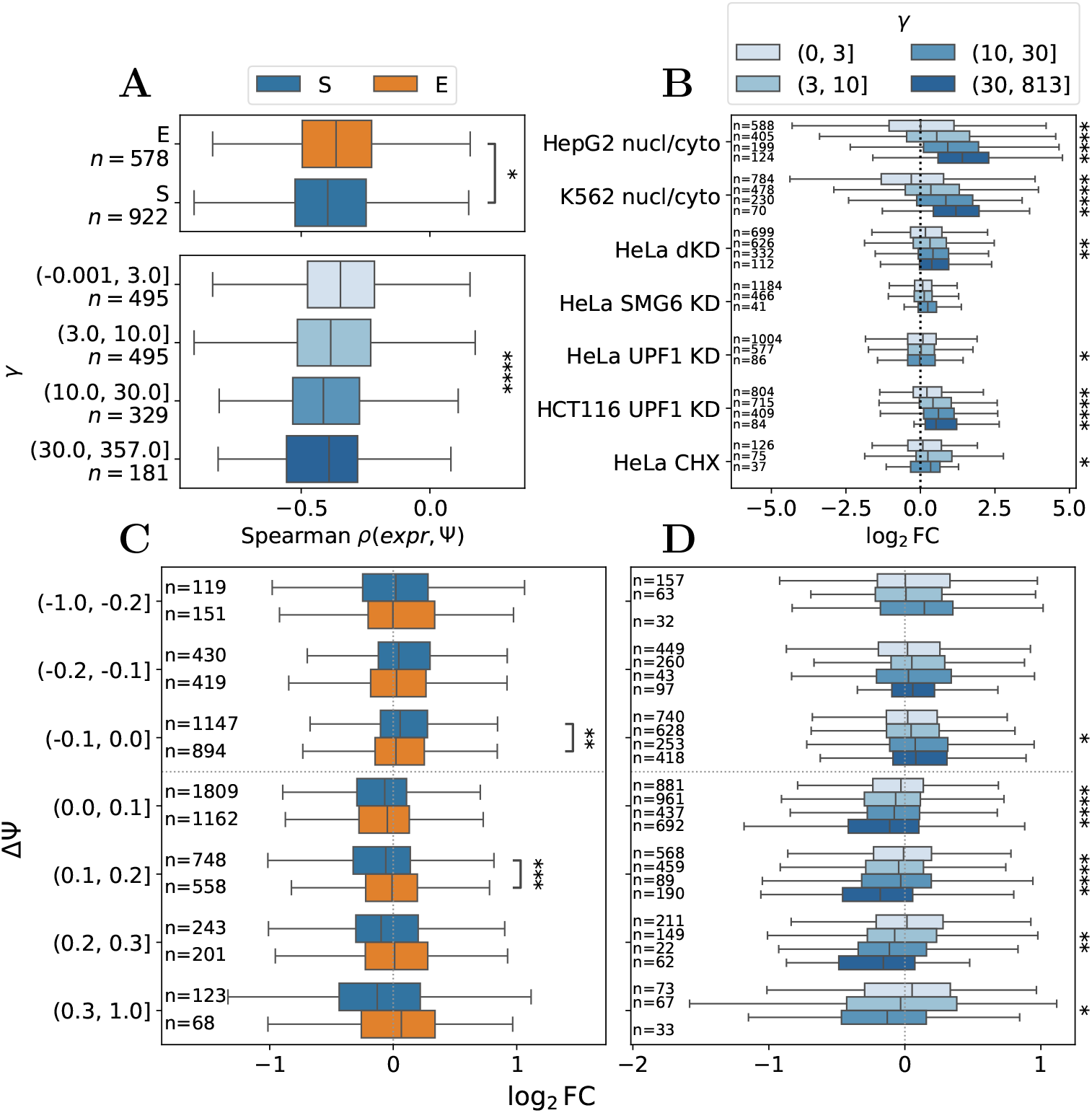
Event-specific relative decay rates predict functional consequences of unproductive splicing. **(A)** Spearman correlation coefficients between exon inclusion levels (Ψ) and host gene expression (*expr*) across GTEx samples. Top panel: NMD-sensitive (S) and NMD-edaving (E) classification. Bottom panel: subdivision into four *γ* groups. **(B)** Host gene expression changes (log_2_ *FC*) upon NMD inhibition for genes subdivided into four *γ* groups. **(C,D)** Host gene expression changes (log_2_ *FC*) in perturbation experiments stratified by ΔΨ and S/E classification (C) or ΔΨ and subdivision into four *γ* groups (D). Asterisks denote significance levels based on Mann–Whitney tests for pairwise comparisons and linear regression analysis for *γ* trends (****, *p* < 0.0001; ***, *p* < 0.001; **, *p* < 0.01; *, *p* < 0.05, after Bonferroni correction).

To determine whether *γ* values predict transcriptional responses to NMD inhibition, we assigned each gene the highest *γ* value among its AS events and classified them into four groups by increasing *γ*. Genes with higher *γ* values generally showed larger increases in expression following NMD inhibition than genes with lower *γ* values (Figure 4B). This relationship was most pronounced in nuclear–cytoplasmic fractionation experiments, with significant differences also observed after SMG6/SMG7 codepletion and UPF1 KD in HCT116 cells.

We next investigated the extent to which event-specific relative degradation rates capture the relationship between gene expression and splicing changes in NMD isoforms and how they compare with the widely used ΔΨ metric, which quantifies changes in exon inclusion between conditions. To this end, we selected RNA-seq datasets from ENCODE consortium profiling RNA-binding protein perturbations, many of which target splicing factors and therefore encompass a broad range of ΔΨ values [30]. For each perturbation, we quantified changes in exon inclusion (ΔΨ) and host gene expression (log_2_FC), subdividing genes into groups according to NMD-sensitive vs. NMD-evading attribution of their associated splicing events and controlling for the magnitude and the direction of ΔΨ.

For ΔΨ > 0, the decrease in gene expression level was consistently larger for NMD-sensitive than for NMD-evading events, while for ΔΨ < 0, the effects were opposite and less statistically significant (Figure 4C). A similar trend was observed when genes were subdivided into four *γ* groups, with progressively stronger decrease in the expression level for ΔΨ > 0 and only weak and statistically insignificant opposite trends for ΔΨ < 0 (Figure 4D). Thus, comparable splicing changes (ΔΨ) produced markedly different transcriptional outcomes for NMD-sensitive and NMD-evading events.

These results demonstrate that the functional impact of differential splicing events generating NMD isoforms cannot be inferred from ΔΨ alone, but must also account for the parameter *γ* of NMD-mediated transcript degradation. Incorporating event-specific *γ* estimates therefore provides a more accurate interpretation of the relationship between AS and gene expression.

### Experimental validation

To evaluate the model under biologically relevant conditions, we selected six NMD-sensitive AS events for RT-qPCR validation in PC3 prostate adenocarcinoma cells. In these cells, experiments were conducted with and without NMD inhibition by CHX treatment, and relative changes in transcript isoforms abundances were quantified using RT-qPCR (Figure 5).

**Figure 5:**
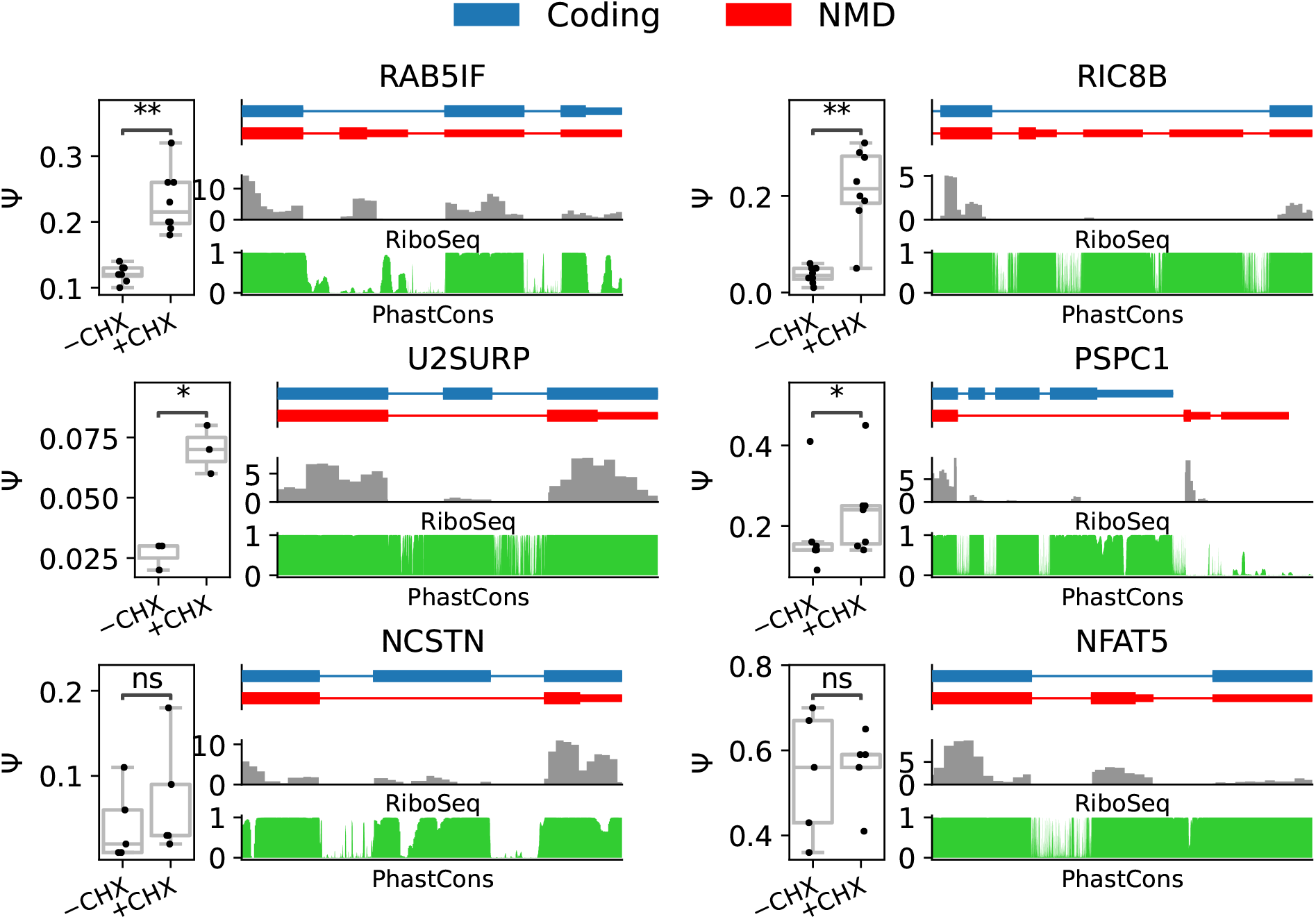
Experimental validation of selected NMD-sensitive events by RT–qPCR in PC3 cells. Each subplot corresponds to one event and is labeled by the host gene name. Within each sub-plot, the left panel shows experimentally measured Ψ values in control and CHX-treated cells. The ideograms display (top-down) two alternative splice isoforms (NMD isoform in red), ribosome profiling coverage under NMD inhibition (gray), and PhastCons30 score (green). Transcript identifiers and genomic coordinates are listed in Table S5. Asterisks indicate significance levels of paired *t*-tests comparing control and CHX-treated samples after Bonferroni correction (****, *p* < 0.0001; ***, *p* < 0.001; **, *p* < 0.01; *, *p* < 0.05).

CHX treatment significantly increased the abundance of the NMD isoforms of *RAB5IF*, *RIC8B*, *U2SURP*, and *PSPC1*, confirming their sensitivity to NMD-mediated degradation. The essential exon in *NCSTN* displayed the same tendency, although the increase was not statistically significant. Ribosome profiling performed under NMD inhibition provided complementary evidence that the poison exons of *RAB5IF* and *PSPC1* could be translated, with pronounced ribosome footprint coverage terminating abruptly at the PTC. This pattern is consistent with active translation of the poison exon under NMD inhibition followed by translation termination at the PTC. Notably, *PSPC1* and *U2SURP* lacked Ribo-seq coverage over exons included in the protein-coding isoform under NMD inactivation but regained it under conditions with intact NMD (Figure S9).

In contrast, the AS event in *NFAT5*, predicted as NMD-sensitive based on SMG6/SMG7 double KD experiments in HeLa cells, did not show a measurable increase in NMD isoform abundance upon CHX treatment in the PC3 cell line. Ribosome profiling revealed a markedly different translational pattern for this gene. Whereas ribosome occupancy in RAB5IF and PSPC1 dropped immediately after the PTC, ribosome footprints in NFAT5 remained detectable beyond the stop codon (Figures 5 and S9). Its estimated readthrough efficiency exceeded that of 77% and 82% of NMD-sensitive events in two independent ribosome profiling datasets (see Methods). However, because ribosome profiling and RT-qPCR were performed in different cell lines, these observations alone do not establish that translational readthrough accounts for the lack of detectable NMD sensitivity in PC3 cells.

To further contextualize these findings, we assessed the inferred NMD sensitivity of the *NFAT5* splicing event using RNA-seq data from UPF1 KD experiments in HCT116 cells matching the conditions used for ribosome profiling [29]. In comparison to other events examined, *NFAT5* showed only a modest response to UPF1 depletion, with Ψ = 0.4 in control and Ψ = 0.6 in UPF1-depleted cells, resulting in a low relative decay rate *γ*≃2 (Figure S10). Consequently, this event was classified as NMD-evading in the HCT116 cell line, in accordance with the readthrough observed by ribosome profiling in these cells, whereas in the HeLa cells under SMG6/SMG7 double KD it was classified as NMD-sensitive. Together, the results suggest a potential role for translational readthrough in modulating the fate of the *NFAT5* NMD isoform, although further experiments in matching cellular contexts would be required to establish a causal link.

## Discussion

The coupling of AS with NMD constitutes an important mechanism of post-transcriptional gene regulation [54, 55, 56]. Current estimates suggest that up to 15% of transcripts produced from protein-coding loci are subject to NMD, and that the impact of AS on the transcriptome is mediated to a greater extent through the production of NMD isoforms than through proteome diversification [56]. Our study aims at addressing quantitative interpretation of RNA-seq measurements of these isoforms, whose steady-state abundances are shaped by selective RNA degradation.

The analytical framework developed here links the observed Ψ values to the underlying splicing rates *r* through a family of functions parameterized by the relative degradation rate parameter *γ*. Linear, nonlinear, and derivative-at-zero fitting approaches revealed substantial discrepancies among the resulting *γ* estimates which, in part, arise from distinct behavior of the regression models across different regions of the (Ψ, *r*) space. At the same time, the apparent heterogeneity of the data, with some events showing dramatic increase upon NMD inhibition and others exhibiting little or no evidence of differential degradation, indicates that the relationship between Ψ and *r* cannot be adequately described by a single universal value of *γ*.

Gaussian mixture modeling confirmed the existence of two large classes of AS events differing in NMD sensitivity, NMD-sensitive and NMD-evading, with markedly different average degradation rates and considerable variability of *γ* values within each class. On the one hand, this variability reflects the underlying biological reality in which transcripts harboring PTCs are not equally sensitive to NMD. The degradation efficiency depends on multiple features, including stop codon position, RNA secondary structure, translation efficiency, and interactions with RNA-binding proteins [57, 43, 46]. Moreover, NMD itself is not a single mechanism but rather a collection of partially overlapping pathways involving distinct combinations of core factors and degradation routes [58].

On the other hand, part of the observed variability arises from differences between cell lines and experimental protocols, which target distinct stages of the NMD pathway and vary in the degree to which they inhibit its activity. Consistently, codepletion of SMG6 and SMG7 exerted the strongest effect on NMD, whereas depletions of single factors, indirect suppression of NMD through CHX treatment, and nuclear–cytoplasmic fractionation resulted in more moderate effects. A remarkable example of cell type-specific response is a poison exon in the *NFAT5* gene, which consistently exhibited high *γ* values across all analyzed RNA-seq datasets except UPF1 KD in HCT116 cells, but did not respond to CHX treatment in PC3 cells and showed evidence of translational readthrough in HCT116 cells. This is consistent with the notion that NMD sensitivity may vary between cellular contexts and that some poison exons can undergo conditional NMD escape [59, 60, 61]. However, despite these discrepancies, the classification into NMD-sensitive and NMD-evading events remained broadly consistent across experiments, indicating that a core set of AS events that trigger NMD can be reproducibly identified.

It should be noted, however, that many validated NMD-sensitive events exhibit strong evolutionary conservation not only within the poison exon itself but also in the flanking intronic regions (Figure 5). This pattern is a well-established hallmark of AS–NMD-mediated regulation, particularly in genes encoding splicing factors [21, 62]. Strong and protracted conserved regions were observed in *RIC8B*, *U2SURP*, and also *NFAT5*, supporting the existence of functional regulatory elements in all these genes.

Another important finding is that accounting for event-specific relative degradation rates improves the interpretation of RNA-seq data by clarifying how AS relates to both NMD sensitivity and gene expression. The difference in exon inclusion levels, ΔΨ, is a conventional metric of differential splicing that is commonly used in two distinct contexts. In NMD-inactivation experiments, ΔΨ can quantify the splicing response to NMD inhibition. However, it does not distinguish NMD-sensitive from NMD-evading events, which can exhibit similar ΔΨ values. That is, two AS events with the same ΔΨ = 0.2, one changing from Ψ = 0.1 to Ψ = 0.3 and the other from Ψ = 0.4 to Ψ = 0.6, may produce NMD isoforms with markedly different degradation rates (Figure 6). This is exemplified by the case of *NFAT5*, where despite a relatively large ΔΨ = 0.6 − 0.4 = 0.2 in UPF1-depleted HCT116 cells, this event was classified as NMD-evading, underscoring that a substantial splicing response to NMD inhibition does not necessarily imply strong NMD sensitivity. In this context, *γ* provides a more direct measure of NMD sensitivity than ΔΨ alone.

**Figure 6:**
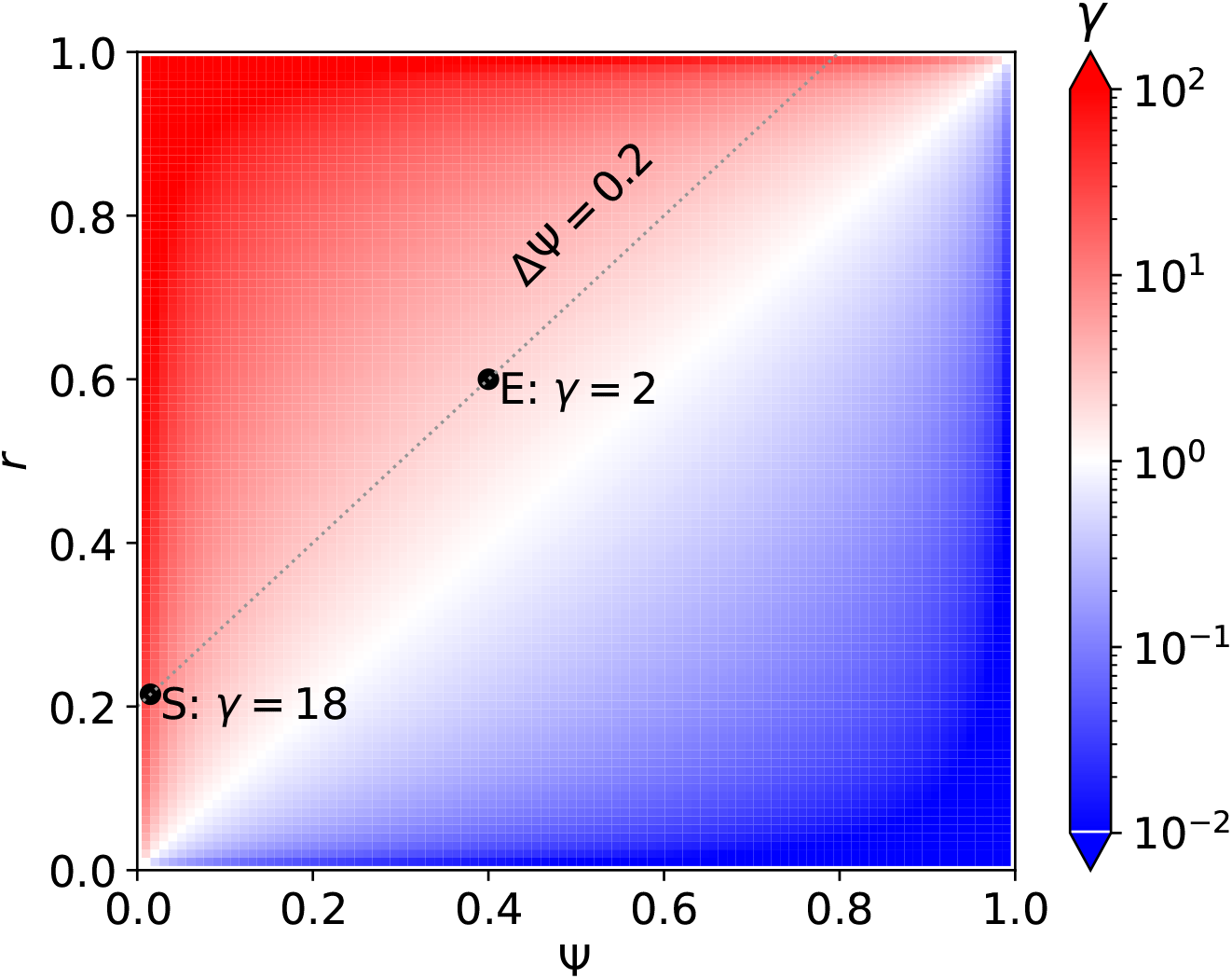
The splicing response to NMD inhibition, measured as ΔΨ = *r* −Ψ, does not uniquely determine NMD sensitivity. The heatmap shows *γ* as a function of Ψ (x-axis) and *r* (y-axis). The dotted line indicates ΔΨ = 0.2. Events with the same ΔΨ can have markedly different *γ* values depending on where in the dotted line they are located and, thus, differ in their sensitivity to NMD, as illustrated by the two highlighted points.

The second context is differential splicing under conditions in which NMD activity is not experimentally altered. Here, ΔΨ measures the magnitude of splicing changes, whereas *γ* helps explain its consequences for gene expression: for the same ΔΨ, events with higher *γ* tend to be associated with larger changes in gene expression (Figure 4C,D). Thus, *γ* does not replace ΔΨ as a measure of differential splicing, but rather captures how the relative degradation of NMD isoforms modulates the effect of splicing changes on gene expression.

The kinetic model in Figure 1A is deliberately simplified, as it considers only two splicing outcomes: NMD and protein-coding transcript isoforms. In reality, most genes undergo multiple AS events, resulting in a spectrum of transcript-specific degradation rates rather than a single kinetic parameter. This complexity can be partially accommodated by redefining the Ψ and *r* variables in terms of the combined abundance of all NMD isoforms relative to the total abundance of all isoforms produced by a given locus, thereby reducing the spectrum of kinetic constants to a single effective value. Without this reduction, the model becomes substantially more complex and is no longer amenable to the simple analytical framework presented here. However, it remains unclear whether further increasing model complexity will yield substantial benefits, as benchmarking gene sets with single versus multiple AS events produces similar parameter estimates (Figure S11).

A curious reader may still ask what range of *γ* values is reasonable. Beyond the observation that it depends on the experimental setting and cellular context, one can confidently state that a reasonable estimate for most NMD-generating AS events is on the order of 10, i.e., NMD isoforms decay at least one order of magnitude faster than protein-coding isoforms. At the same time, its value depends on the specific AS event, cellular context, and overall NMD efficiency, and may act as a regulatory tuning parameter in diverse biological processes.

## Conclusion

This work lays the theoretical foundation for relating splicing metrics derived from RNA-seq data to the underlying splicing rates of NMD-sensitive transcripts. It also has substantial practical significance, as it enables the estimation of corrected AS rates from the vast body of RNA-seq data accumulated in the literature to date, thereby improving downstream analyses and the biological interpretation of alternative splicing.

## Authors’ contributions

LZ and DP designed the study; LZ and MV performed data analysis; AK and DS performed the experiments; LZ and DP wrote the first draft of the manuscript. All authors edited the final version of the manuscript.

## Acknowledgments

The authors cordially thank Prof. Sergei A. Spirin for valuable comments on the manuscript.

## Competing interests

The authors declare no competing interests.

## Funding

This work was supported by the Russian Science Foundation grant 22-14-00330-P.

## Code availability

All scripts used to analyze data and generate figures are available through a ZENODO reposi-tory [40].

## Notes

### Competing Interest Statement

The authors have declared no competing interest.

### Summary of Updates

Additions: RNA-seq data matching Ribo-seq; RF classifier for NMD escape covariates; additional analysis for NMD escape in NFAT5

