## Supplementary material for "Estimation of splicing metrics for NMD-sensitive transcripts"

October 15, 2026

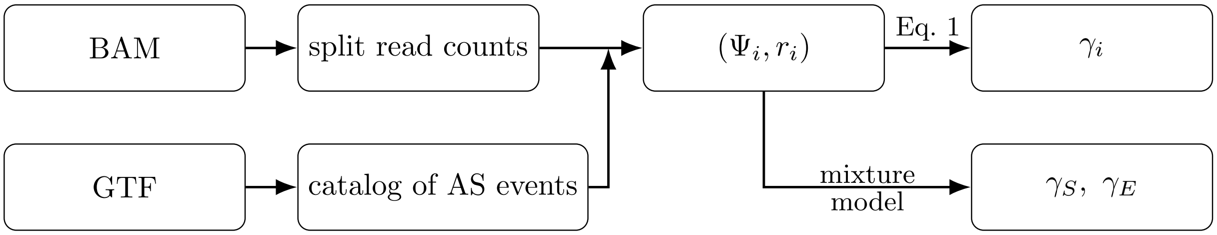

Figure S1: A schematic overview of the analysis pipeline. Split-read counts are extracted from BAM files and assigned to AS events defined by the catalog constructed from the annotation (GTF). The corresponding counts are used to calculate splicing ratios $\Psi_i$ and $r_i$ under control and NMD-inhibited conditions, respectively. These ratios are then used to estimate event-specific relative transcript decay rates $\gamma_i$. A Gaussian mixture model is fitted to the splicing data to estimate the relative decay rates of the NMD-sensitive and NMD-evading components, $\gamma_S$ and $\gamma_E$, respectively.

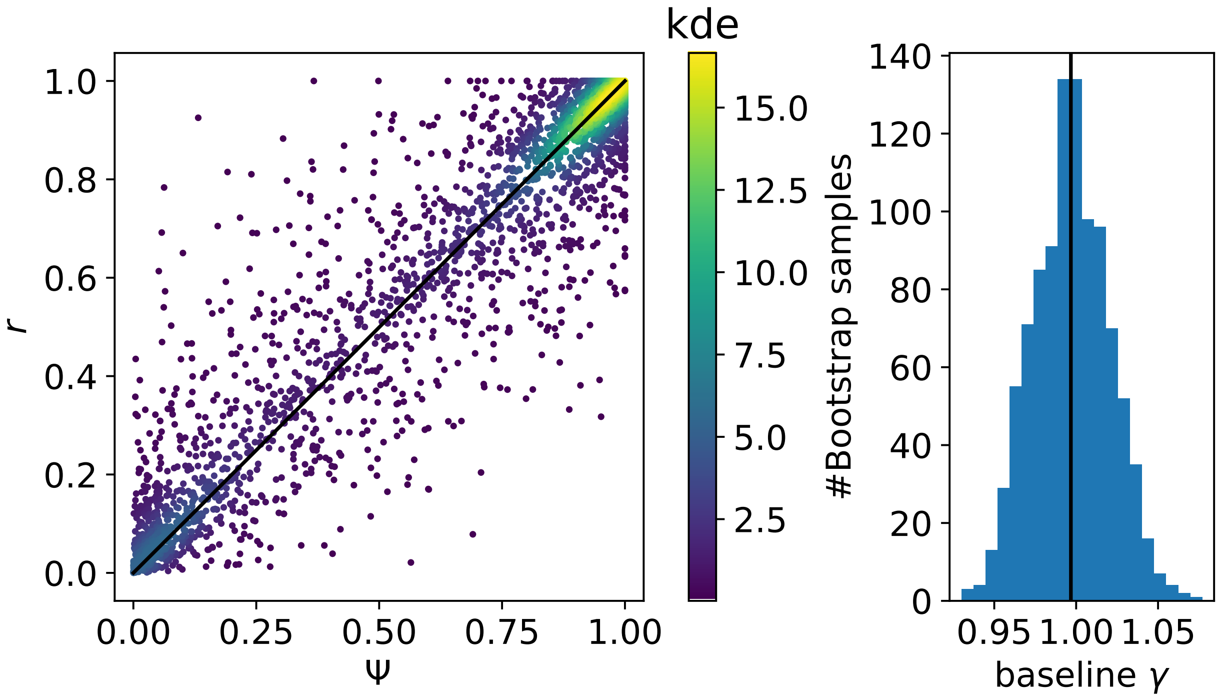

Figure S2: Estimation of the baseline decay parameter $\gamma_E$. Left: Density plot of $r$ versus $\Psi$ for a set of AS events generating only protein-coding isoforms. Colors represent probability density (kde). Right: Distribution of $\gamma_E$ estimates obtained from bootstrap sampling of the same set of events.

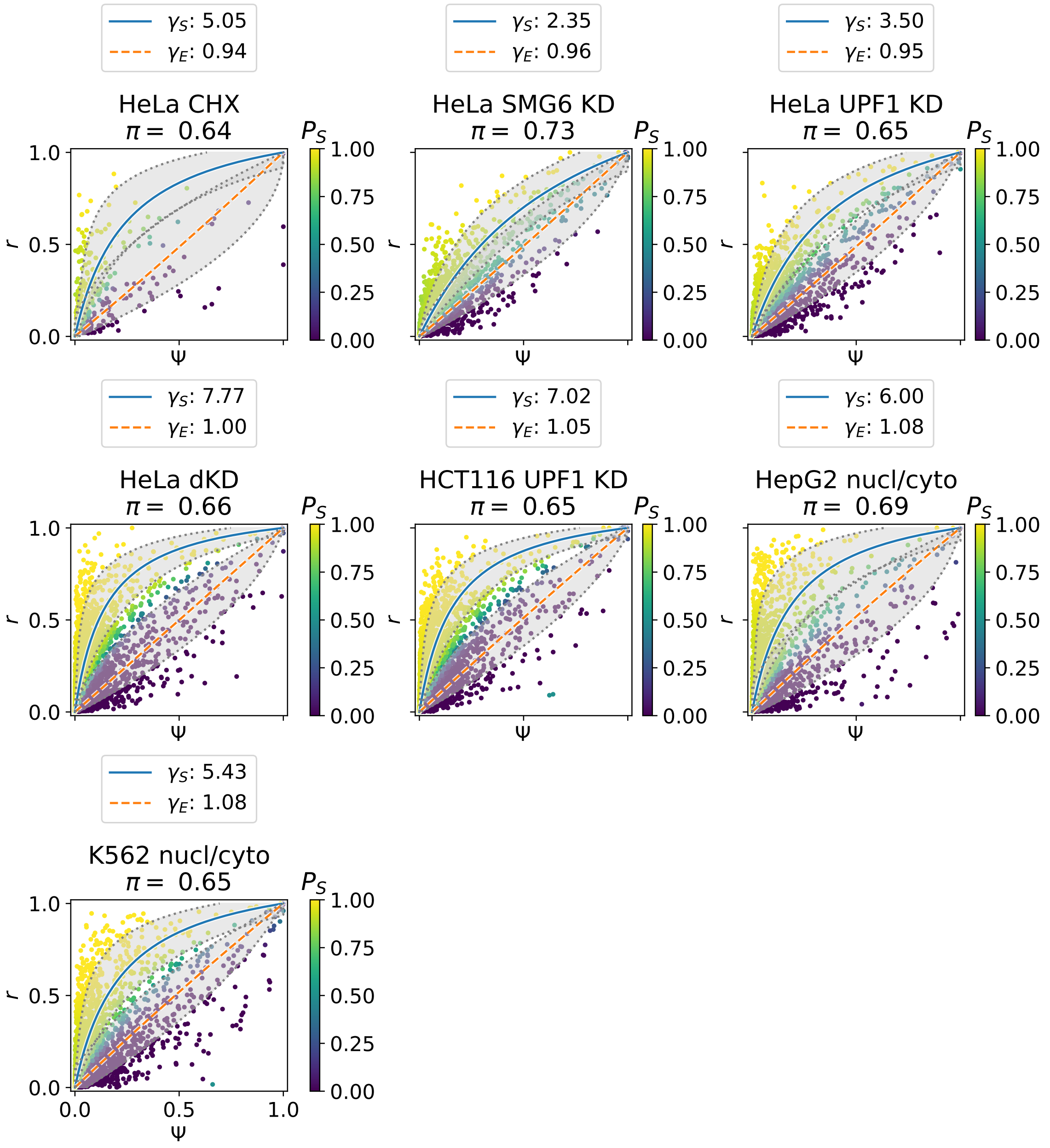

Figure S3: Two-component Gaussian mixture model on the full set of NMD-generating AS events applied to seven independent RNA-seq datasets: HeLa CHX treatment, SMG6 knockdown, UPF1 knockdowns in HeLa and HCT116 cells, SMG6/SMG7 codepletion, and nuclear–cytoplasmic fractionation in HepG2 and K562 cells. The rest of the legend is as in Figure 3C.

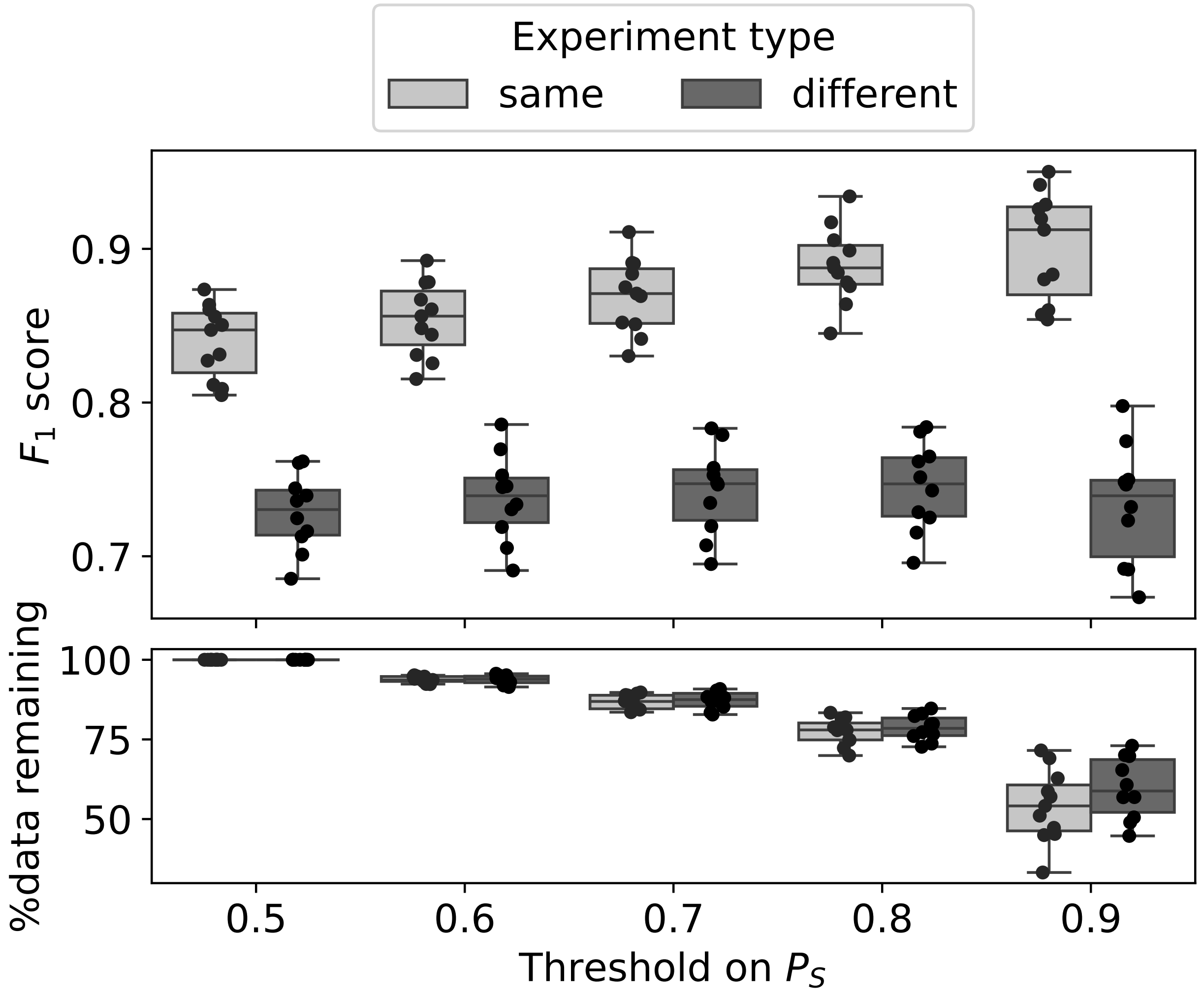

Figure S4: Robustness of AS event classification to the probability threshold. AS events were classified as NMD-sensitive if $P(S \mid \Psi, r) > t$ and as NMD-evading if $P(S \mid \Psi, r) \leq 1 - t$, with events satisfying $1 - t < P(S \mid \Psi, r) \leq t$ left unclassified. Upper panel: $F_1$-scores for pairwise comparisons; each point represents a pairwise comparison. Colors indicate comparisons within the same experimental group (NMD-inhibition experiments or nuclear–cytosolic fractionation) or comparisons between these groups. Lower panel: the percentage of AS events retained for classification after excluding unclassified AS events.

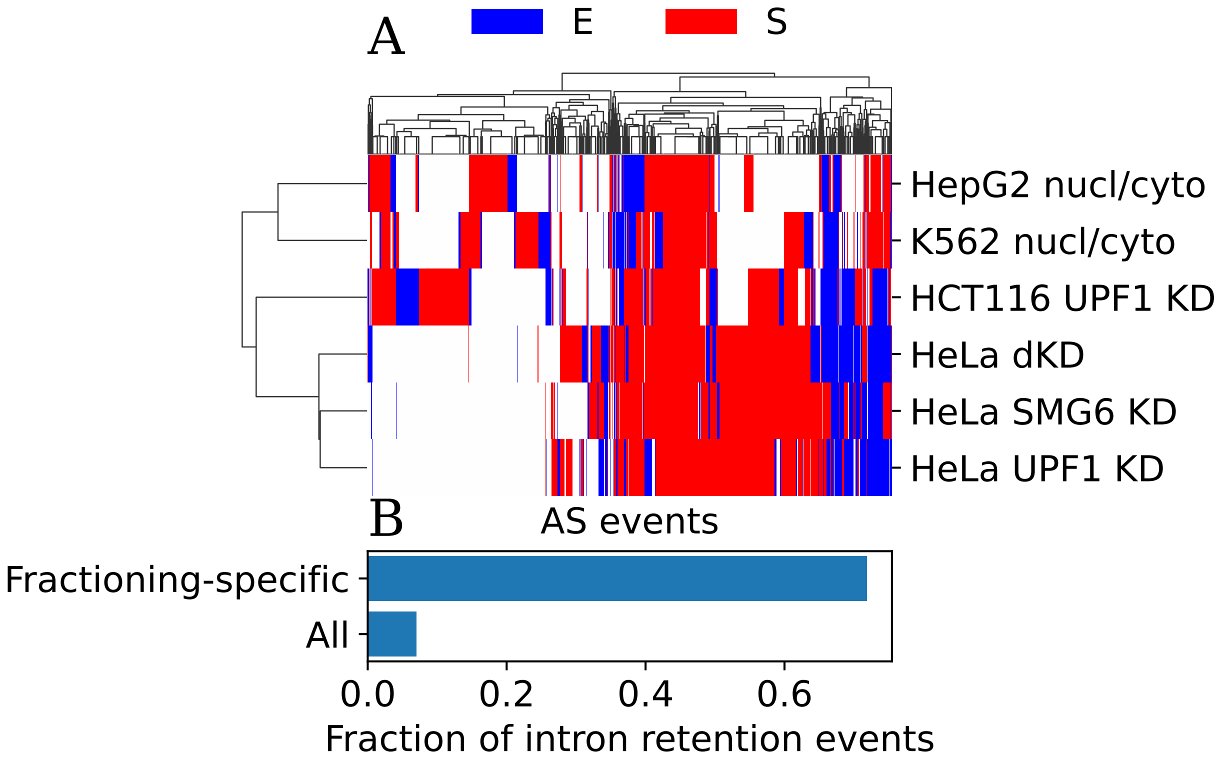

Figure S5: NMD-sensitivity classifications across AS events and experimental datasets. **(A)** Clustered heatmap showing event classifications across experiments, with each cell indicating NMD-sensitive (S, red) or NMD-evading (E, blue) status. **(B)** The proportion of intron retention events among all analyzed events and among events classified as NMD-sensitive specifically in nuclear–cytosolic fractionation, but not in other experiments.

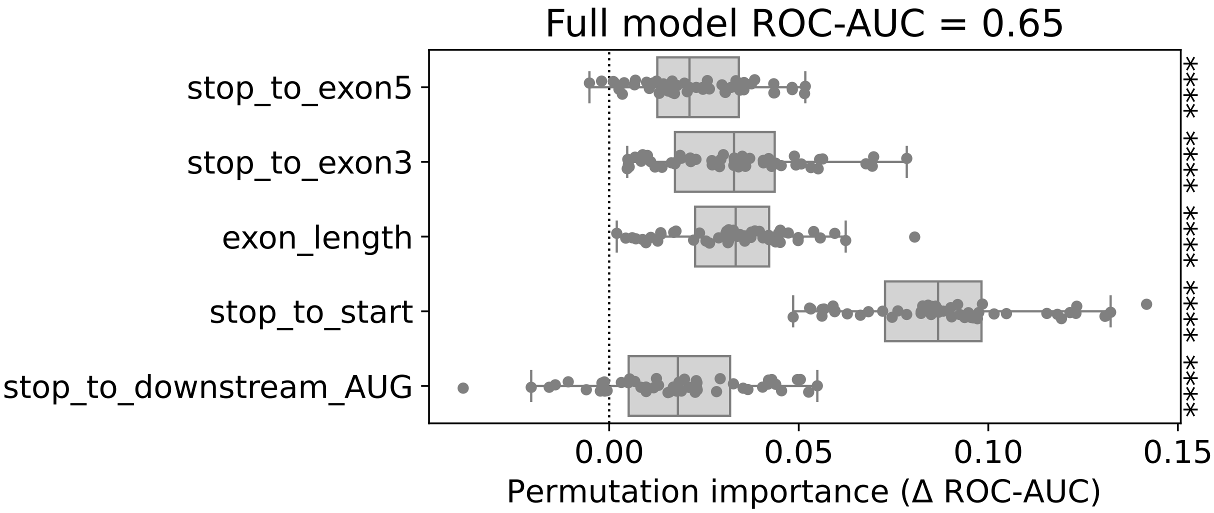

Figure S6: Feature importance for random forest classification of NMD-sensitive and NMD-evading events. Feature importance was quantified as the decrease in ROC-AUC when a feature was permuted relative to the full model trained on the original data. Model performance was evaluated using stratified five-fold cross-validation. For each fold, one full model was trained and its ROC-AUC was calculated; each feature was then permuted independently 10 times, and ROC-AUC of the resulting models were compared with that of the full model. The boxplots show the resulting importance estimates across 50 permutations per feature (10 permutations in each of five folds).

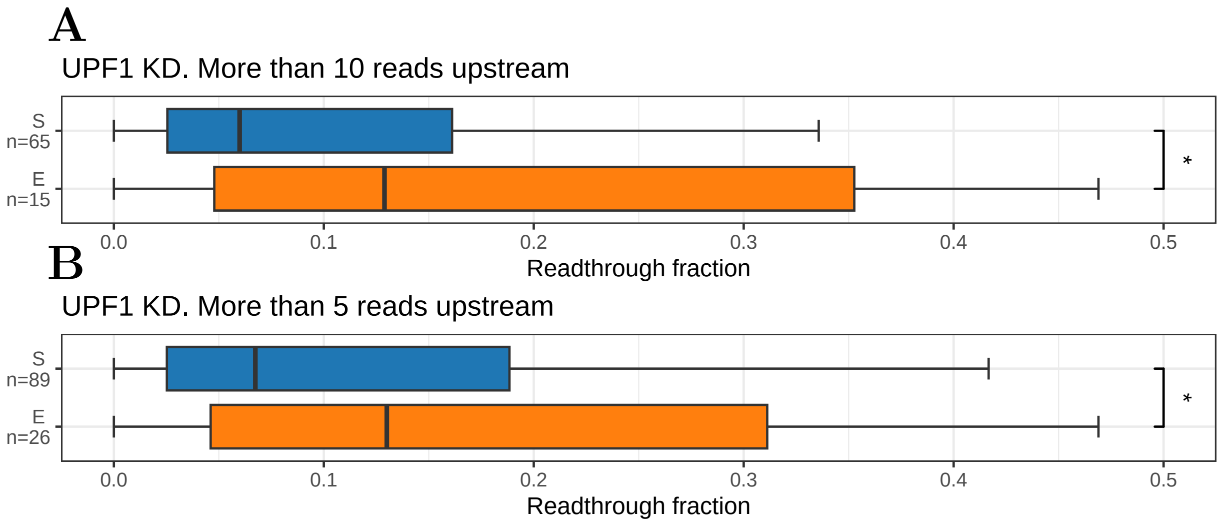

Figure S7: Readthrough fractions in NMD-sensitive and NMD-evading poison exons after filtering for minimum Ribo-seq coverage upstream of the PTC. Readthrough fraction was calculated as in Figure 3H. AS events with more than 10 (A) or more than 5 (B) reads in the region upstream of the PTC are shown.

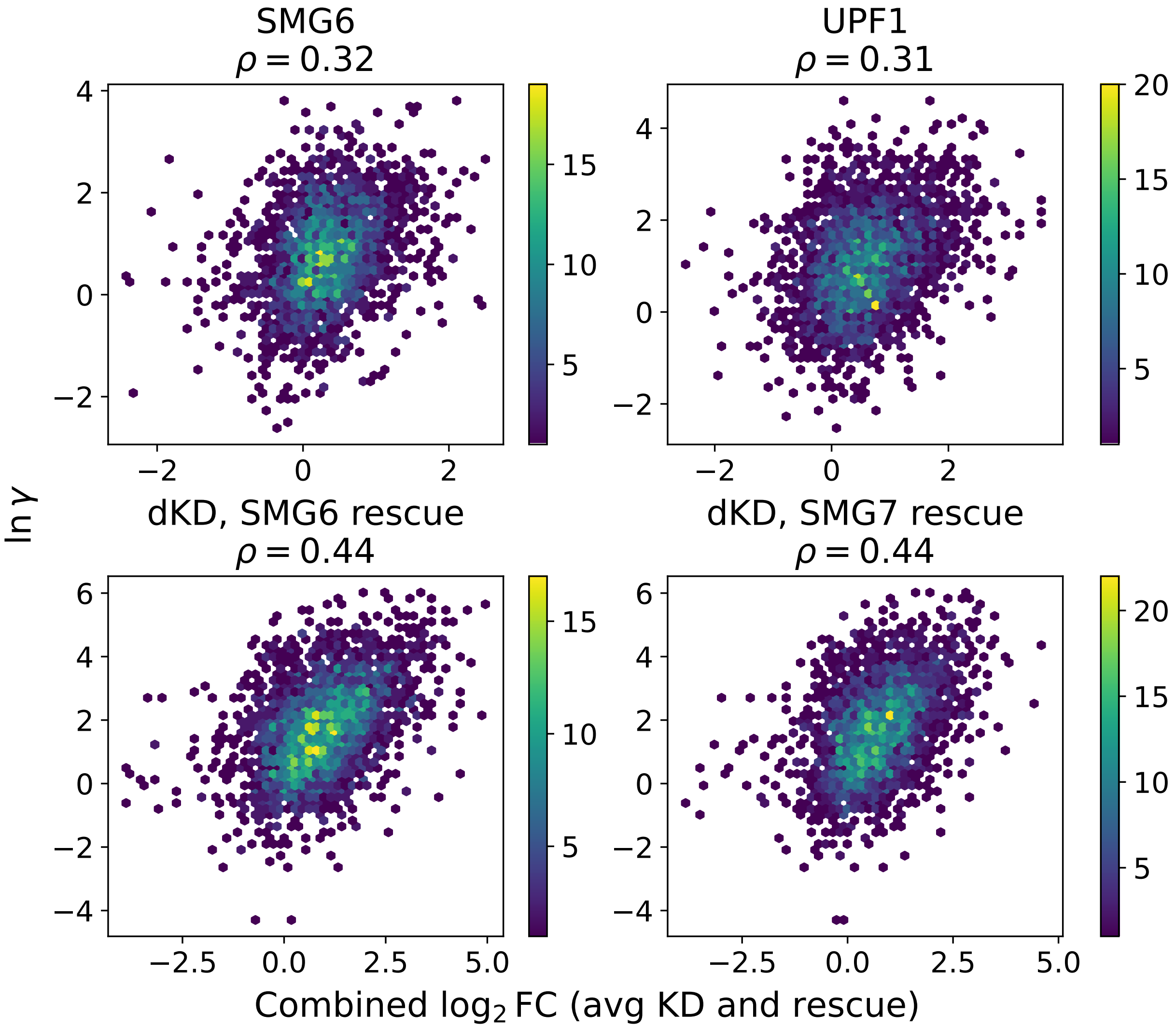

Figure S8: Correlation between event-specific relative decay rates and independent transcript-level measures of NMD efficiency. Hexbin plots show log-transformed event-specific $\gamma_i$ values against the combined NMD response $\log_2$ fold-change scores from Colombo et al. for SMG6 knockdown, UPF1 knockdown, and SMG6/SMG7 double knockdown with two rescue conditions. Color scales indicate the number of observations within each hexagonal bin.

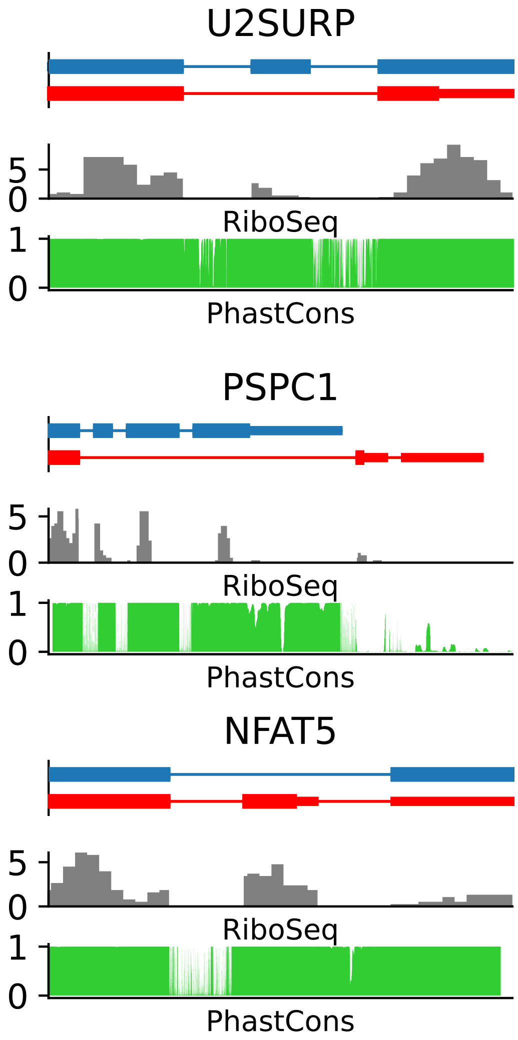

Figure S9: Ribosome profiling coverage in *U2SURP*, *PSPC1*, and *NFAT5* without NMD inhibition. The rest of the legend is as in Figure 5.

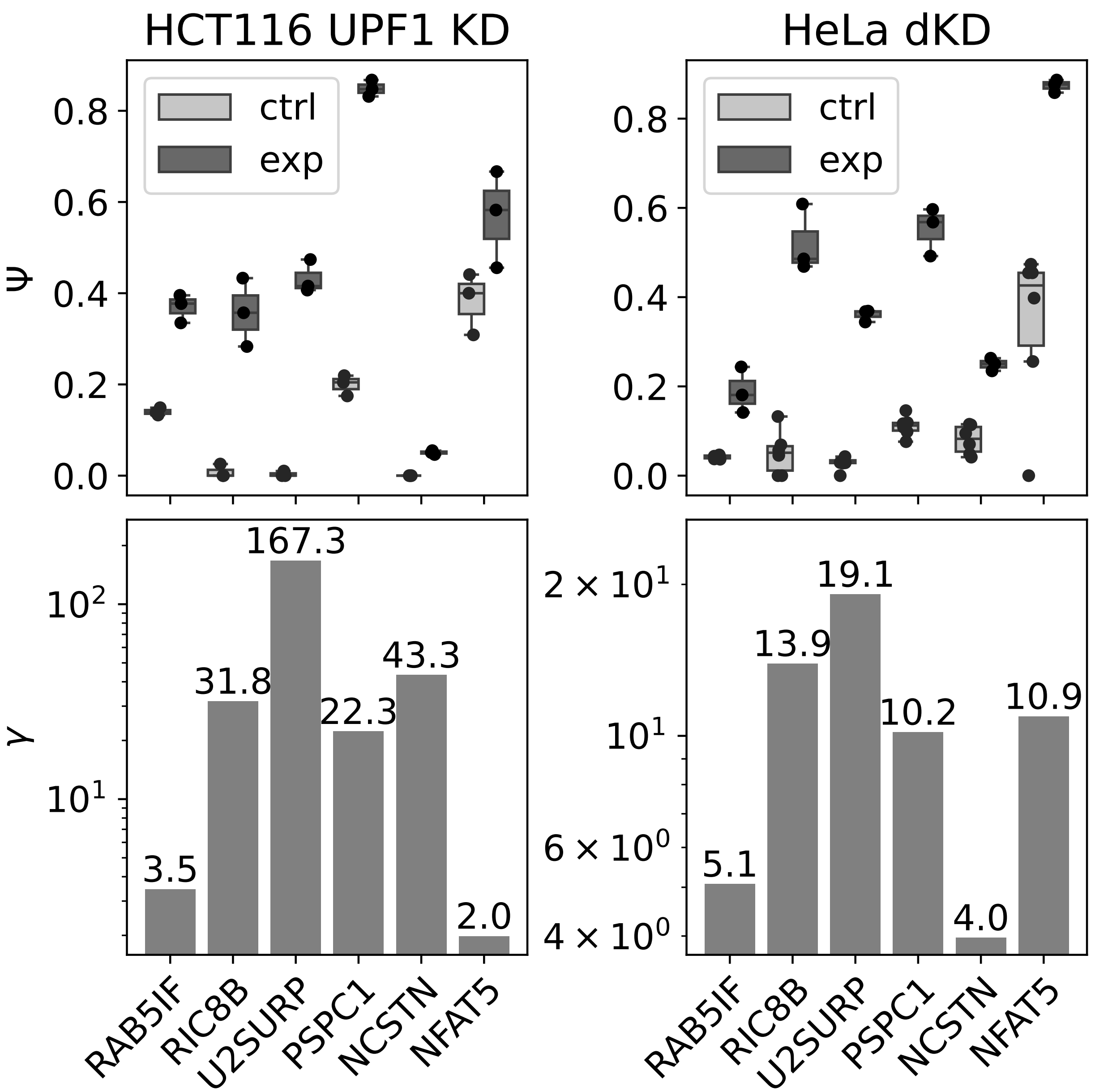

Figure S10: Splicing ratios (Ψ) and estimated relative decay rates ($\gamma$) of selected AS events in HCT116 (left) and HeLa cells (right). Upper panels show Ψ values in the control and knockdown conditions; lower panels show the corresponding $\gamma$ estimates.

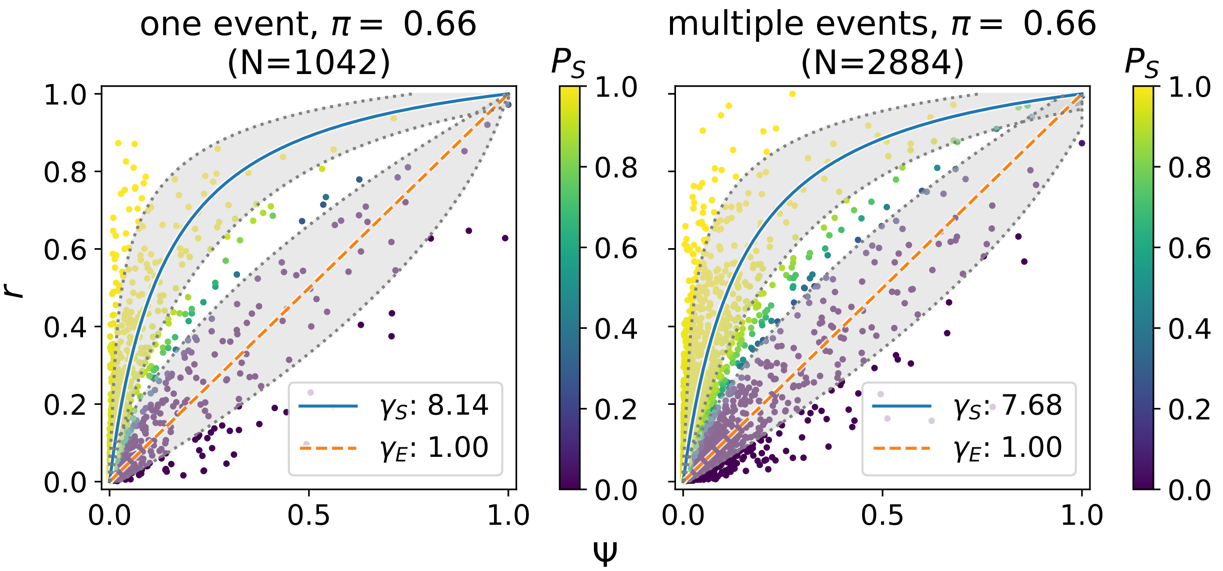

Figure S11: Gaussian mixture models fitted to SMG6/SMG7 co-depletion RNA-seq data in genes with single vs. multiple AS events. Left panel: mixture models fitted to AS in genes with a single AS event. Right panel: mixture models fitted to AS in genes with multiple AS events. The rest of the legend is as in Figure 3C.

| Primer | Sequence |
|---|---|
| RAB5IF_F | GATGTGATCTACTGGTTCCGAC |
| RAB5IF_R | CAGGCAGAATCCTGCTATTCC |
| RAB5IF_3aR | CAATGGCCTTTCCCTGCTATT |
| U2SURP_F | ATGCTCCTATGTTACCGCCA |
| U2SURP_13R | TGGCTTGCGACAGAGT |
| U2SURP_R | GGCGAGCAAATTCCTTCTCAA |
| RIC8B_F | ACTTTCCAGAGAGGAGTTGCT |
| RIC8B_R | TGTCAGAGCTGGTCTCTTCG |
| RIC8B_10aF | GATGACTAAGACCCTGTCATTA |
| RIC8B_11R | CGCTTGCCACTTAGCTTGAT |
| PSPC1_F | ACATGAAGAGGAGCATCGGC |
| PSPC1_R | CCTGTTCTCTATTTTCCATGTAGTTTG |
| PSPC1_utrR | TTATCACCATTTTCCATGTAGTTTGG |
| NCSTN_F | GCCAGAGTTTGCTCACTGC |
| NCSTN_6R | TCTTGATAGCACTGCTTGATGAC |
| NCSTN_R | CAGACGATTTCTGCTTGATGAC |
| NFAT5_F | GTCAGACAAGCGGTGGTGAG |
| NFAT5_R | TGAAGAAGCATCAGCAGCAAC |
| NFAT5_4aR | AGAGGCAAATCCAGCAGCAAC |

Table S1: qPCR primer sequences.

| Tissue | Number of samples |
|---|---|
| Artery - Aorta | 246 |
| Artery - Coronary | 140 |
| Artery - Tibial | 357 |
| Brain - Amygdala | 81 |
| Brain - Anterior cingulate cortex (BA24) | 99 |
| Brain - Caudate (basal ganglia) | 134 |
| Brain - Cerebellar Hemisphere | 115 |
| Brain - Cerebellum | 144 |
| Brain - Cortex | 132 |
| Brain - Frontal Cortex (BA9) | 117 |
| Brain - Hippocampus | 103 |
| Brain - Hypothalamus | 104 |
| Brain - Nucleus accumbens (basal ganglia) | 123 |
| Brain - Putamen (basal ganglia) | 103 |
| Brain - Spinal cord (cervical c-1) | 76 |
| Brain - Substantia nigra | 71 |
| Breast - Mammary Tissue | 218 |
| Colon - Sigmoid | 173 |
| Fallopian Tube | 7 |
| Heart - Atrial Appendage | 218 |
| Heart - Left Ventricle | 267 |
| Lung | 372 |
| Minor Salivary Gland | 70 |
| Muscle - Skeletal | 470 |
| Nerve - Tibial | 334 |
| Ovary | 108 |
| Pancreas | 192 |
| Pituitary | 124 |
| Testis | 198 |

Table S2: GTEx tissue samples.

| Experiment | Sample type | SRA run |
|---|---|---|
| HeLa NMD KD | ctrl | SRR4081222 |
| HeLa NMD KD | ctrl | SRR4081223 |
| HeLa NMD KD | ctrl | SRR4081224 |
| HeLa NMD KD | ctrl | SRR4081237 |
| HeLa NMD KD | ctrl | SRR4081238 |
| HeLa NMD KD | ctrl | SRR4081239 |
| HeLa NMD KD | SMG6 KD | SRR4081231 |
| HeLa NMD KD | SMG6 KD | SRR4081232 |
| HeLa NMD KD | SMG6 KD | SRR4081233 |
| HeLa NMD KD | UPF1 KD | SRR4081225 |
| HeLa NMD KD | UPF1 KD | SRR4081226 |
| HeLa NMD KD | UPF1 KD | SRR4081227 |
| HeLa NMD KD | SMG6+SMG7 dKD | SRR4081246 |
| HeLa NMD KD | SMG6+SMG7 dKD | SRR4081247 |
| HeLa NMD KD | SMG6+SMG7 dKD | SRR4081248 |
| HeLa CHX | ctrl | SRR8443314 |
| HeLa CHX | ctrl | SRR8443315 |
| HeLa CHX | CHX | SRR8443318 |
| HeLa CHX | CHX | SRR8443319 |
| HepG2 nucl/cyto | cyto | SRR4422482 |
| HepG2 nucl/cyto | cyto | SRR5048088 |
| HepG2 nucl/cyto | cyto | SRR4422481 |
| HepG2 nucl/cyto | cyto | SRR5048087 |
| HepG2 nucl/cyto | nucl | SRR5048082 |
| HepG2 nucl/cyto | nucl | SRR4422653 |
| HepG2 nucl/cyto | nucl | SRR5048081 |
| HepG2 nucl/cyto | nucl | SRR4422652 |
| K562 nucl/cyto | cyto | SRR5048067 |
| K562 nucl/cyto | cyto | SRR4421723 |
| K562 nucl/cyto | cyto | SRR5210711 |
| K562 nucl/cyto | cyto | SRR5048068 |
| K562 nucl/cyto | cyto | SRR4421722 |
| K562 nucl/cyto | nucl | SRR5048097 |
| K562 nucl/cyto | nucl | SRR4421919 |
| K562 nucl/cyto | nucl | SRR5048098 |
| K562 nucl/cyto | nucl | SRR4421920 |
| HCT116 UPF1 KD | ctrl | E-MTAB-13788: 191508.bam |
| HCT116 UPF1 KD | ctrl | E-MTAB-13788: 191510.bam |
| HCT116 UPF1 KD | ctrl | E-MTAB-13788: 191512.bam |
| HCT116 UPF1 KD | UPF1 KD | E-MTAB-13788: 191532.bam |
| HCT116 UPF1 KD | UPF1 KD | E-MTAB-13788: 191534.bam |
| HCT116 UPF1 KD | UPF1 KD | E-MTAB-13788: 191536.bam |

Table S3: Accession numbers of RNA-seq experiments.

| Gene name | Source | Type | Chr | Start | End | Strand | PMID |
|---|---|---|---|---|---|---|---|
| ACTR5 | chess | A3SS | chr20 | 38765518 | 38766257 | + | 24567363 |
| DCLK2 | ensembl | PE | chr4 | 150249684 | 150256020 | + | 36912101 |
| FUS | ensembl | EE | chr16 | 31185179 | 31188325 | + | 21358643, 24204307 |
| HNRNPA2B1 | chess | IR | chr7 | 26190460 | 26190993 | - | 20946641, 23863836 |
| HNRNPD | ensembl | A3SS | chr4 | 82355001 | 82355304 | - | 17000771, 29263134 |
| HNRNPL | ensembl | PE | chr19 | 38840559 | 38843842 | - | 18073345, 19124611, 27808105 |
| HNRNPLL | chess | PE | chr2 | 38577532 | 38581913 | - | 18073345, 19124611 |
| IQGAP1 | ensembl | PE | chr15 | 90482281 | 90483361 | + | 36912101 |
| MDM4 | chess | complex | chr1 | 204532246 | 204538209 | + | 26595814 |
| PTBP1 | chess | EE | chr19 | 806556 | 808360 | + | 14731397, 17679092, 27808105 |
| PTBP2 | ensembl | EE | chr1 | 96804939 | 96806866 | + | 17606642, 17679092, 21723171, 27808105 |
| RBM39 | ensembl | PE | chr20 | 35739017 | 35740824 | - | 30916337 |
| RPL3 | ensembl | A3SS | chr22 | 39317021 | 39317461 | - | 16254077 |
| RPS3 | ensembl | A5SS | chr11 | 75400777 | 75401640 | + | 30916337 |
| SF1 | ensembl | EE | chr11 | 64770408 | 64776498 | - | 24637117 |
| SFPQ | chess | complex | chr1 | 35177399 | 35187001 | - | 30916337 |
| SNRNP70 | ensembl | PE | chr19 | 49101471 | 49104634 | + | 23389473 |
| SNRPB | ensembl | PE | chr20 | 2465819 | 2467607 | - | 18443041, 21325135 |
| SRSF1 | chess | complex | chr17 | 58004001 | 58004923 | - | 33176162 |
| SRSF10 | ensembl | complex | chr1 | 23978273 | 23978653 | - | 17361132, 33176162 |
| SRSF11 | ensembl | complex | chr1 | 70228555 | 70231095 | + | 17361132, 33176162 |
| SRSF11 | ensembl | PE | chr1 | 70228555 | 70232268 | + | 33176162 |
| SRSF3 | ensembl | PE | chr6 | 36598983 | 36601152 | + | 17361132, 22436691, 26416554, 26704980 |
| SRSF4 | ensembl | complex | chr1 | 29160517 | 29181646 | - | 33176162 |
| SRSF6 | ensembl | PE | chr20 | 43458509 | 43459771 | + | 33176162 |
| SRSF7 | ensembl | PE | chr2 | 38748653 | 38749529 | - | 22436691, 24637117, 30916337, 33176162 |
| SRSF8 | chess | IR | chr11 | 95068183 | 95068655 | + | 33176162 |
| SRSF9 | ensembl | PE | chr12 | 120464122 | 120465627 | - | 17361132, 33176162 |
| TIA1 | ensembl | PE | chr2 | 70224629 | 70227735 | - | 24637117 |
| TRA2A | ensembl | complex | chr7 | 23522432 | 23531789 | - | 24637117, 33176162 |
| TRA2B | ensembl | PE | chr3 | 185926734 | 185937825 | - | 33176162 |
| U2AF1 | ensembl | PE | chr21 | 43100519 | 43101366 | - | 30916337 |

Table S4: List of validated AS events. Event types: PE – poison exon, EE – essential exon, A5SS – alternative 5'-splice site. A3SS – alternative 3'-splice site, IR – intron retention.

| Gene name | NMD transcript ID | Coding transcript ID | Coordinates/Link |
|---|---|---|---|
| RAB5IF | ENST00000483815 | ENST00000344795 | chr20:36607818-36612010 |
| RIC8B | ENST00000462949 | ENST00000392837 | chr12:106860412-106885904 |
| U2SURP | ENST00000488497 | ENST00000473835 | chr3:143023064-143027149 |
| PSPC1 | ENST00000492741 | ENST00000619300 | chr13:19674984-19730239 |
| NCSTN | ENST00000459963 | ENST00000699553 | chr1:160350250-160352887 |
| NFAT5 | ENST00000393742 | ENST00000349945 | chr16:69626528-69647028 |

Table S5: Transcript identifiers and genomic locations of AS events in Figure 5.
